# Uncertainty in denoising of MRSI using low-rank methods

**DOI:** 10.1101/2021.05.15.444311

**Authors:** William T Clarke, Mark Chiew

**Affiliations:** Wellcome Centre for Integrative Neuroimaging, FMRIB, Nuffield Department of Clinical Neurosciences, University of Oxford, Oxford, United Kingdom

**Author notes:** To whom correspondence should be addressed **Corresponding author contact details:** Dr William T. Clarke Wellcome Centre for Integrative Neuroimaging, FMRIB University of Oxford Level 0 John Radcliffe Hospital Oxford OX3 9DU UK.

**Keywords:** Spectroscopy, MRS, MRSI, low rank, denoising

## Abstract

**Purpose:** Low-rank denoising of MRSI data results in an apparent increase in spectral SNR. However, it is not clear if this translates to a lower uncertainty in metabolite concentrations after spectroscopic fitting. Estimation of the true uncertainty after denoising is desirable for downstream analysis in spectroscopy.

In this work the uncertainty reduction from low-rank denoising methods based on spatio-temporal separability and linear predictability in MRSI are assessed. A new method for estimating metabolite concentration uncertainty after denoising is proposed. Finally, automatic rank threshold selection methods are assessed in simulated low SNR regimes.

**Methods:** Assessment of denoising methods is conducted using Monte Carlo simulation of proton MRSI data, and by reproducibility of repeated in vivo acquisitions in five subjects.

**Results:** In simulated and in vivo data, spatio-temporal based denoising is shown to reduce the concentration uncertainty, but linear prediction denoising increases uncertainty. Uncertainty estimates provided by fitting algorithms after denoising consistently under-estimate actual metabolite uncertainty. However, the proposed uncertainty estimation, based on an analytical expression for entry-wise variance after denoising, is more accurate. Finally, it is shown automated rank threshold selection using Marchenko-Pastur distribution can bias the data in low SNR conditions. An alternative soft-thresholding function is proposed.

**Conclusion:** Low-rank denoising methods based on spatio-temporal separability do reduce uncertainty in MRS(I) data. However, thorough assessment is needed as assessment by SNR measured from residual baseline noise is insufficient given the presence of non-uniform variance. It is also important to select the right rank thresholding method in low SNR cases.

## Introduction

Increasing the signal-to-noise ratio (SNR) of magnetic resonance spectroscopic imaging (MRSI) allows for faster acquisitions, more reliable quantification, or higher resolution acquisitions. One way SNR can be increased is by reducing the noise variance through computational post-processing, i.e., by ‘denoising’. Low-rank denoising methods achieve this in spectroscopic imaging data either by exploiting the linear predictability, or the spatio-temporal separability of the spectroscopic data, or both (1). Low-rank denoising is a data-driven technique and does not incorporate prior knowledge or physical models of the data. These methods have also recently been applied to MR imaging techniques that utilise an additional dimension of encoding, such diffusion encoding direction in diffusion-weighted MRI (2,3), and time in functional MRI (4). Low-rank models have also been applied directly in the reconstruction of fast MRSI acquisitions (5,6). The application of low-rank models to these techniques aims to exploit signal correlations across these encoding dimensions.

High levels of apparent denoising are consistently achieved by denoising algorithms in MRS with additional encoding dimensions (7,8), or MRSI (1,9,10). However, it is not clear whether there is an overall reduction in uncertainty of final dynamic model parameters (11), or metabolite concentrations (the typical output of MRSI). Denoising will lower the apparent noise in any given spectrum but can introduce systematic model-based errors affecting the bias and variance of the output, ultimately resulting in lower reproducibility and higher mean squared error. Furthermore, as spectroscopic signals are typically converted to metabolite concentrations by fitting of an explicit spectroscopic model to the data (12), it is not clear whether denoising prior to fitting is statistically advantageous. Finally, uncertainty estimation of the fitted metabolite concentrations by fitting packages typically assumes a uniform spectral (or time-domain) noise profile. Usually a frequency-independent noise variance is estimated from a signal-free region or noise pre-scans. After low-rank denoising, the residual noise cannot be assumed to be independent and identically distributed (complex) Gaussian noise. Therefore, uncertainty estimates cannot be trusted without validation. For the same reason, SNR is an inadequate metric for the evaluation of denoising algorithms.

Therefore, in first part of this work we assess actual uncertainty reduction achieved by low-rank algorithms based on linear predictability (LP) of the time-domain signal (13), spatio-temporal (ST) separability (14), and a combination of the two: using the Low Rank Approximations method (LORA) (1). We do this using Monte Carlo (MC) simulations of a toy problem, in simulated ^1^H-MRSI data, and finally by assessing reproducibility in vivo ^1^H-MRSI of the human brain. We additionally assess a new variance propagation method for data truncated by singular value decomposition for accurate estimation of the fitting uncertainties in denoised MRSI data (15,16).

In MRSI, multiple metabolite signals are present in each voxel, the amplitudes of each signal are expected to vary across space, primarily driven by changes in metabolite concentration. Where there is pathologically driven change, metabolite concentrations can vary dramatically across short distances. To maximise noise variance reduction, low-rank methods aim to explain the data using as few rank components as possible. However, using few components may not accurately capture the full variation of rapidly changing metabolite concentrations. Automated parameter selection methods such as those based on Stein's Unbiased Risk Estimate (SURE) minimise the mean squared error (MSE) of the denoised data (17,18), whilst Marchenko-Pastur (MP) based methods aim to separate noise from signal by estimating which components arise from the pure noise matrix (2). But minimisation of spectral data MSE is not equivalent to minimisation of fitted concentration MSE. In very noisy data, or in data where the target signal variance is on the order of the noise variance, methods which minimise data MSE are unlikely to preserve true signal variation.

Therefore, in the final part of this work, in ST denoising of simulated data, we assess the effect of different rank estimation methods and a different thresholding operation.

## Theory

### Low-rank denoising methods

In this work truncation to a fixed rank is achieved using the singular value decomposition. Given an observation model

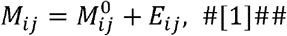

the Eckart-Young-Mirsky theorem states the best rank-r approximation to 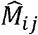 can be derived from a truncated singular value decomposition retaining only the *r* highest singular values

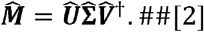

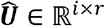 and 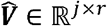 contain the truncated left and right singular vectors of 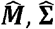 is a truncated diagonal matrix of singular values, and † is the conjugate transpose. The noise term in [1], *E_ij_*, may either be zero-mean i.i.d. complex Gaussian (for ST denoising), or otherwise structured (for LP denoising).

#### Linear predictability (LP) denoising

LP denoising aims to exploit the low-rankness of a Hankel matrix formed from the single-voxel time domain signal (13). This low-rankness arises from the sparsity of the equivalent spectral information.

This method is applied voxel-wise to the MRSI data, with no data shared across voxels. A Hankel matrix is formed from the time-domain data of a single voxel *s*

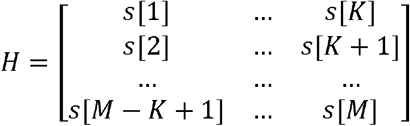

And the denoised Hankel matrix 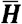 is formed by minimising

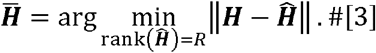

The denoised time-domain data is then reformed from the first row and last column of 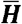. Effectively “softening” the low-rank filtering by not enforcing the Hankel structure in Equation [3] (1).

#### Spatio-temporal (ST) denoising

ST denoising exploits the partial separability of spatio-temporal modes due to correlated spectral information across space. I.e. that as in Equation 2 of Reference (1) the signal can be seen as L^th^ -order separable between the spatial basis *c_l_* (*r*) and the spectral basis *ψ*_1_(*f*)

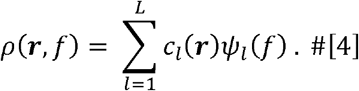

Equation 4 may equivalently be written in terms of basis in the temporal domain

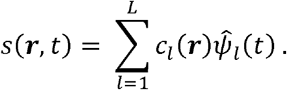

To perform ST denoising a Casorati matrix is formed

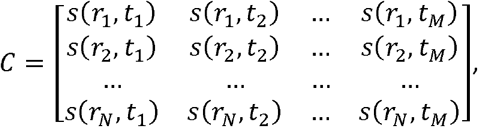

and the denoised matrix 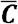 is formed by minimising

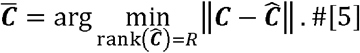

#### LORA

The LORA method combines ST and LP denoising sequentially, applying ST to all voxels simultaneously before applying LP denoising voxel-wise.

#### Patch based methods (ST and LORA)

ST and LORA can be applied globally to all voxels in a dataset, or locally in an overlapping patch-wise manner. In the local method, ST denoising is applied within each patch, and the signals from voxels belonging to multiple (overlapping) patches are averaged. Both methods may be applied to a restricted range of voxels identified by a mask. The patch is characterised by a 3D patch size and a stride parameter, which dictates the amount of patch overlap.

### Uncertainty of low-rank denoised data

According to Chen *et al*, under moderate-to-high SNR conditions, the variance of an element 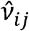 of ‘denoised’ matrix 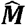 is

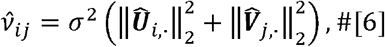

where *σ*^2^ is the variance of the i.i.d. noise of the observed data (15).

Song *et al* extended the above framework to incorporate patch-based low-rank denoising methods (16). The variance of elements having undergone patch denoising and averaging is given by Equation 20 of Reference (16). This extension of Chen *et al’s* method incorporates calculation of variance cross terms in the presence of mutually shared information between patches.

### Uncertainty of metabolite concentrations from denoised data

The variance of the denoised data may be non-uniform in the time-domain (Figure 1), and there may be significant covariance between the denoised time-domain data points. When this is the case the conventional estimation of metabolite concentration uncertainty by using a signal-free region to estimate noise variance is insufficient. However, propagation of non-uniform and correlated variance through the non-linear fitting process analytically is difficult. Therefore, in this work a bootstrapping approach is proposed. Repeated fits are made of each voxel’s data perturbed with complex correlated noise, created using the estimated variance and covariance of the denoised data. The off-diagonal elements of the covariance of the time-domain data are approximated as with the full covariance matrix estimated by combination with the element-wise variance (Equation 6)

**Figure 1.**
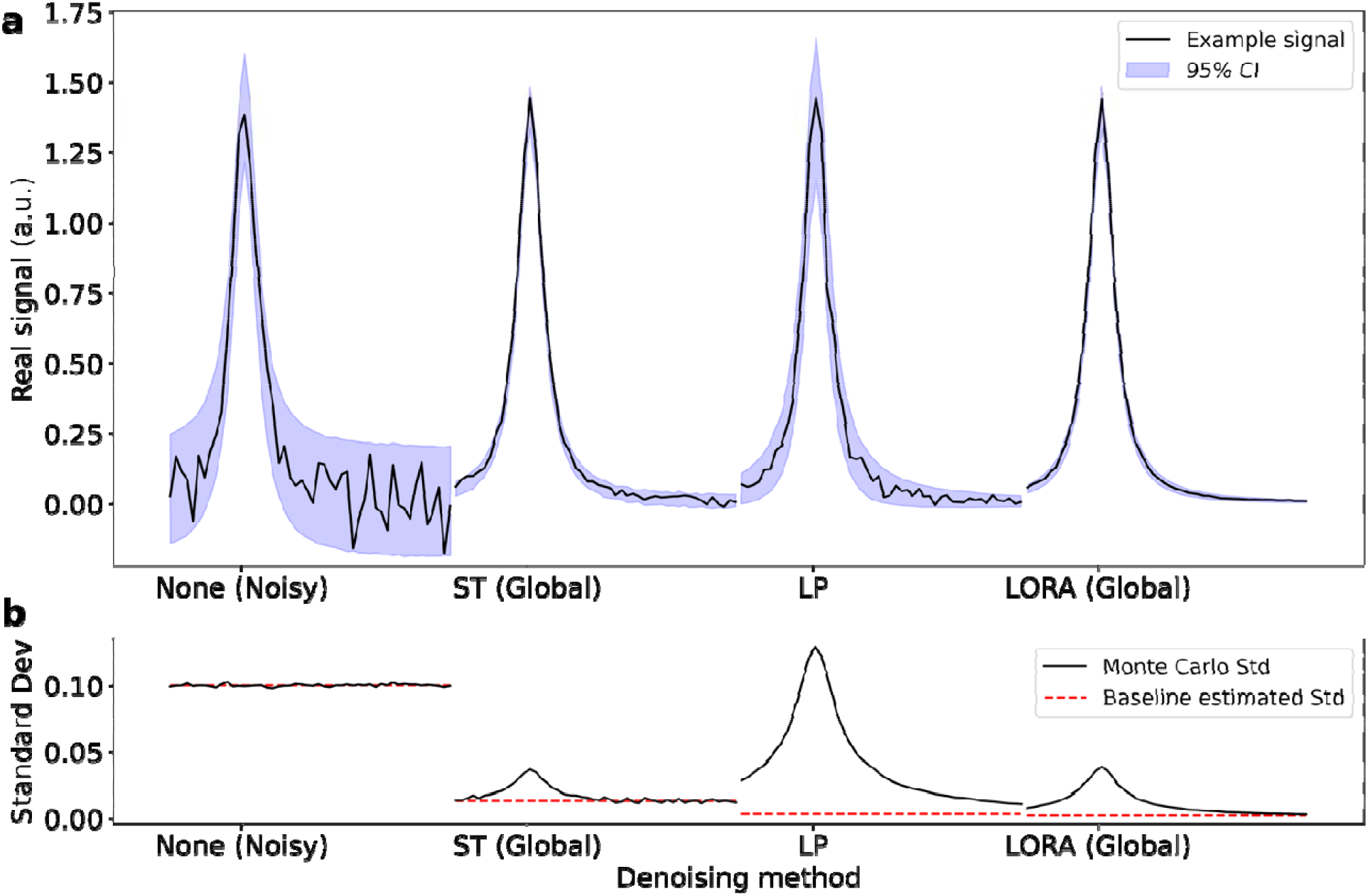
**a** Single peak after application of different low-rank denoising methods (black line) and Monte Carlo 95% confidence intervals (shaded). **b** Monte Carlo standard deviation of the denoised data (black) and the noise level estimated from a signal-free region of baseline (red). Whilst the original noisy data (left) has uniform and high variance the denoised data shows very inhomogeneous variance, with higher values in areas with signal present. Data shown has original noise standard deviation of 0.1. Only a limited frequency range is shown in each panel.

Where is the covariance of time point with time point in voxel *j*, denotes the conjugate transpose operation, and is the variance of the i.i.d. noise of the observed noisy data. The covariance of patch-based methods is estimated as the mean of the covariance of all overlapping patches. A qualitative assessment of this approximation for spectral data is made in the Supporting Information section “Covariance approximation”.

### Automatic rank selection methods: Stein’s Unbiased Risk Estimate and Marchenko-Pastur

Two methods for automatically estimating the rank threshold from the noisy data are assessed in this work. In addition, a different thresholding operation, “soft” or singular value thresholding (SVT), is introduced for one framework.

#### Soft thresholding

Singular value thresholding (SVT) or soft thresholding linearly shrinks the singular values above a threshold otherwise setting values below the threshold to zero. I.e. the SVT estimated denoised matrix is defined as in Candès *et al* (equation 2 in (17))

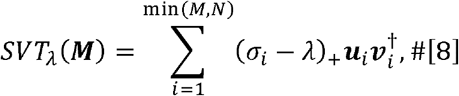

where *x*_+_ = max (*x*, 0), *λ* is the threshold, and *σ_i_* is the i^th^ singular value.

Singular value hard thresholding (SVHT) is implemented for all other cases in this work and may be written,

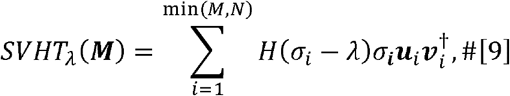

where *H* is the Heaviside function, or equivalently as a function of rank threshold *R*

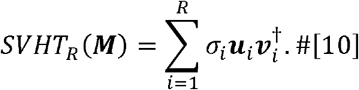

#### SURE

For the model [1], Stein’s Unbiased Risk Estimate (SURE) constructs an unbiased estimate of the risk (mean-square-error) without requiring knowledge of the ground truth. For example, for the SVT function

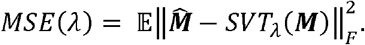

Using SURE it is therefore straightforward to find the singular value threshold *λ* which minimises the MSE of the denoised data. SURE for the (soft) singular value thresholding operation for complex data is given by Equation 7 in reference (17). The equivalent expression for a hard thresholding (SVHT) is given by Equations 2 & 3 in reference (18). The equations as implemented in this work are given in full in the Supporting Information.

#### Marchenko-Pastur

The second method assessed utilizes the upper limit of the Marchenko-Pastur (MP) distribution, which models the distribution of singular values of random matrices, as a singular-value threshold. Introduced by Veraart *et al* (2), this method truncates to zero all eigenvalues below a threshold given by

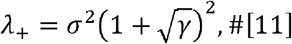

with *γ* = *M* / *N* (the ratio of the dimensions of matrix ***M***) and σ^2^ the variance of the i.i.d. noise of the observed noisy data.

## Methods

All fitting in this work was carried out using version 1.1.2 of FSL-MRS (19). Denoising was carried out using version 0.0.2 of the Python “mrs-denoising-tools” package. Data and analysis code for this paper are available online, please see the data availability statement.

### In vivo ^1^H MRSI data

In vivo ^1^H-MRSI data was acquired from the brains of five healthy subjects (age/weight) at 3 tesla (Magnetom Prisma, Siemens Healthineers, Germany) using a previously reported density-weighted concentric ring (CONCEPT) MRSI sequence with semi-LASER localization (20,21). Reconstructed data formed a 48×48×1 grid at a spatial resolution of 5×5×15 mm^3^. Each subject was scanned with the CONCEPT sequence ten times sequentially. Each separate acquisition was identical, baring a frequency adjustment between acquisitions, and took 4:30 to acquire.

Reconstruction and preprocessing of the data used in-house custom MATLAB scripts designed specifically for the density-weighted sequence. These steps were as follows:

1. Loading and reordering of the non-cartesian k-space data.
2. Regridding with the adjoint 2D non-uniform fast Fourier transform (NUFFT) (22).
3. Coil combination using the wSVD algorithm with weights calculated from the water signal (23).
4. Processing of metabolite-cycled acquisitions to form water-suppressed and unsuppressed data.
5. Phase and frequency correction across the ten averages using cross-correlation (24).
6. Frequency correction across voxels using B_0_ shifts measured from the water reference data.
7. A combined high-SNR average was constructed as the mean of the 10 sequential acquisitions in each subject. This formed data with acquisition time equivalent to 45 minutes.
8. Data was saved in the NIfTI-MRS format with masks derived from hard thresholding of the water reference data, selecting a 16×16×1 block in each subject (25).

All volunteers were recruited and scanned in compliance with local ethical and legal requirements.

### Denoising of uniform single-peak simulation

The performance of the LP, global ST, local ST, and LORA denoising algorithms was evaluated in a simple Monte Carlo test using a single on-resonance Lorentzian peak. Data for this analysis was generated as an 8×8×1 grid of voxels each containing identical signal. The signal was formulated as a decaying exponential with unit amplitude and 10 Hz spectral linewidth (full width at half maximum, fwhm), i.e.

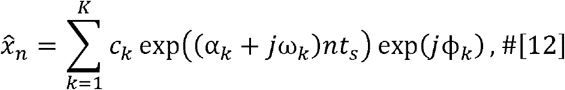

with *K* = 1, *c*_1_ = 1, *ω*_1_ = 1, *α*_1_ = −10*π*, and ϕ_1_ = 0. Fifty instantiations of this data were created with independent and identically distributed complex Gaussian noise at each of six noise levels with standard deviations of 0.5, 0.1, 0.05, 0.01, 0.005, and 0.001.

At each noise level, for each Monte Carlo repetition (separate noise instantiation), data was denoised using the LP, global ST, local ST, and global LORA algorithms. For this explicitly constructed rank = 1 case all algorithm rank thresholds were set to one. All denoised data and original noisy data were fit using non-linear least squares (Scipy ‘curve_fit’ Levenberg-Marquard algorithm (26)) using the same model as was used to generate the data (equation [9]). Each fit produced an estimate of the peak amplitude and the amplitude uncertainty. The Scipy ‘curve_fit’ amplitude uncertainty is derived from the diagonal elements of the covariance matrix, which in turn, is derived from the numerically estimated Jacobian.

For the global and local ST methods the denoised time-domain variance and covariance were estimated using the proposed method (equations [6] & [7]), and subsequently 100 bootstrap fitting repetitions were carried out to estimate the amplitude uncertainty after fitting. The variance and covariance estimated using the proposed method were qualitatively compared to the Monte Carlo estimated variance and covariance.

The “actual uncertainty” of the fitted amplitude for each denoising case and noise level was calculated using the standard deviation across the Monte Carlo repetitions. The “conventional uncertainty” was calculated as the mean of the fitting derived uncertainty. The approximation to the actual uncertainty was calculated as the mean of the bootstrap derived uncertainty. The uncertainty expected for the average of all the voxels in the 8×8×1 grid was also calculated. All uncertainties were then summarized across all noise levels as a ratio to the actual (Monte Carlo derived) uncertainty of the noisy data.

### Evaluation of denoising methods in simulated ^1^H MRSI

Denoising performance and uncertainty estimation were also assessed in realistic simulated ^1^H-MRSI data. Simulated data was constructed from the median concentration, lineshape, frequency shift and noise variance of the five high SNR (45-minute equivalent) in vivo acquisitions. The high SNR acquisitions were fit using the FSL-MRS ‘Newton’ algorithm using a basis set with 17 metabolites (ascorbate [Asc], aspartate [Asp], creatine [Cr], γ-aminobutyric acid [GABA], glucose [Glc], glutamine [Gln], glutamate [Glu], glycerophosphocholine [GPC], glutathione [GSH], myo-inositol [Ins], lactate [Lac], N-acetylaspartate [NAA], N-acetylaspartateglutamate [NAAG], phosphocholine [PCho], phosphocreatine [PCr], scyllo-inositol [scyllo], taurine [Tau]), default FSL-MRS macromolecules and a 3^rd^ -order baseline (19). Only the 256 voxels within the mask of each subject were fitted and used to construct the simulated data. Noiseless simulated data was constructed using the FSL-MRS Voigt lineshape model with baseline and phase parameters set to zero.

From the noiseless simulated data 50 different noise instantiations (MC repetitions) were created for each of five noise levels (Figure 2a). The noise levels were equivalent to 90, 45, 9, 4.5, and 2.25 minutes of scanning. Data was saved in NIfTI-MRS format with a single mask to identify the 256 voxels containing data and water reference scan for consistent metabolite concentration scaling.

**Figure 2.**
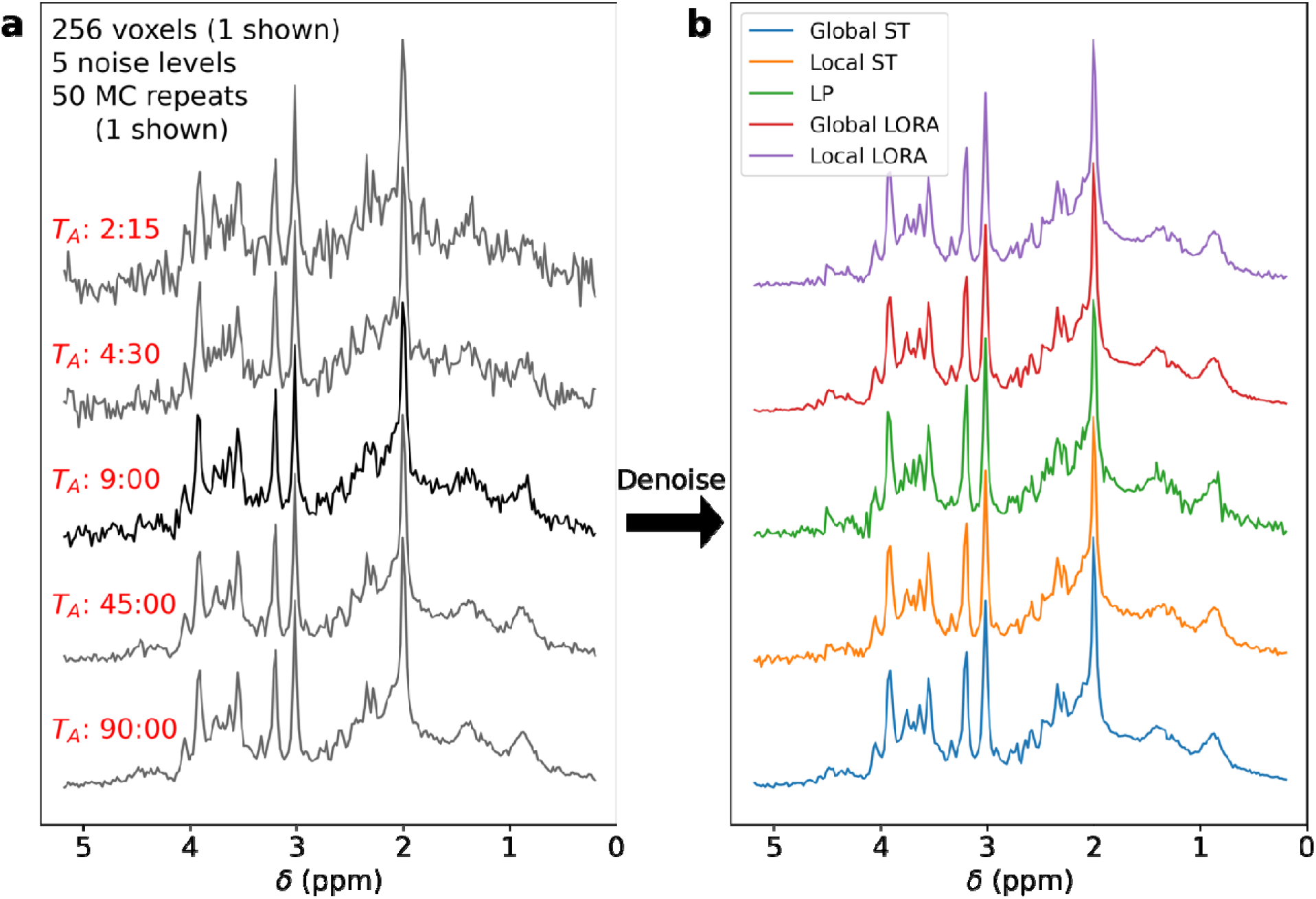
**a** Example synthetic data at each of the five noise levels. In **b,** the 9-minute equivalent data is shown denoised by each of the five methods.

A single noise level (9-minute equivalent) was denoised using LP, global ST, local ST, global LORA, and local LORA (Figure 2b). All other noise levels were denoised using just the global and local ST methods. Rank thresholds were selected using the MP method, except for the LP and second stages of LORA, which used a fixed threshold of 20. Local versions of ST and LORA used a patch size of 4×4×1 voxels with a stride of 1. Denoising (and subsequent fitting) was only applied to the 16×16×1 voxel masked region. Each denoised dataset and the original noisy data was fitted using fsl_mrsi using the parameters and basis set described at the start of this section, but with no baseline (19). The concentrations of highly correlated metabolites were combined (namely Glu+Gln, Glc+Tau, PCho+GPC, NAA+NAAG, Cr+PCr).

Variance and covariance for the ST denoised-data, estimated by the proposed method, was compared with that measured using the 50 MC repetitions. Five of the generated datasets at each noise level underwent bootstrap fitting. For each voxel of the selected datasets, additional correlated noise was added using the estimated covariance matrix before fitting was carried out. This process was repeated 40 times. The reported concentrations and uncertainties are calculated from the mean and standard deviation of the 40 fits.

For analysis, two groups of metabolites were defined: ‘high signal’ metabolites include Glu+Gln, Glc+Tau, PCho+GPC, NAA+NAAG, Cr+PCr and Ins; whilst ‘all unique’ contains all fitted metabolites (combined as described) including the macromolecular peaks.

For the single noise level analysis, mean concentration uncertainties were calculated for all methods and expressed as a fraction of the noisy data uncertainty. The means was calculated across all voxels and metabolites in the respective sets (‘high signal’ and ‘all unique’). RMSE was calculated and normalized using the fitted concentrations of the noiseless synthetic data. RMSE were calculated across all voxels and metabolites in the respective sets (‘high signal’ and ‘all unique’).

For the ST methods applied at all noise levels the MC ‘actual uncertainty’, FSL-MRS ‘conventional uncertainty’, and the bootstrap estimated ‘actual uncertainty’ were calculated for all voxels and metabolites in the respective sets (‘high signal’ and ‘all unique’). These values are expressed as ratios to the MC ‘actual uncertainty’ of the noisy data. A ratio < 1 indicates lower uncertainty, i.e., better performance.

### Reproducibility of in vivo ^1^H MRSI

For each of the five subjects, the 10 sequential in vivo acquisitions were denoised as described above for the simulated data. The resultant denoised spectra, the original noisy spectra, and the combined high SNR (45-minute equivalent) spectra were then fit. Fitting was performed as described above using fsl_mrsi (3^rd^ order baseline, 17 metabolites, default FSL-MRS macromolecules, ‘Newton’ algorithm with Voigt lineshape) (19). Fitting was performed over the 256 masked voxels. Concentrations were references to the unsuppressed water reference scans for each subject, but no relaxation correction was carried out (i.e. T_E_=0 ms, T_R_=15 s). Therefore, metabolite concentrations are expressed in ‘institutional-units’.

The effect of each denoising method was then assessed by comparing the RMSE of the metabolite concentrations of denoised 4.5-minute acquisitions to those from the 45-minute acquisition. Additionally, reproducibility was assessed as the standard deviation of the fitted metabolite concentrations across the 10 repeated acquisitions for each subject. This value was normalized for each metabolite (by the median concentration) and expressed as a ratio to the median of the voxel-wise reproducibility of the original noisy spectra.

### Bias and rank selection in simulated data

To examine the efficacy of the two rank threshold selection methods in low SNR data, synthetic MRSI data was prepared with an explicitly constructed rank-3 Casorati matrix. This was achieved by generating three spectral peaks (at −200, 0 and 300 Hz offsets) with different spatially varying concentrations in an 8×8×1 grid. This numerical phantom is shown in supporting figure S1. Subsequently the phantom was generated with 50 MC repetitions at six different noise levels. The highest noise levels were picked to ensure that the MP and SURE rank threshold estimation techniques underestimated the rank of the data (supporting figure S2).

Each MC repetition in each noise level was denoised using five different approaches to rank threshold selection combined with local ST denoising. The rank threshold was either selected using the MP method, fixed at rank 2, fixed at rank 3, selected using the SURE SVHT algorithm, or the SURE SVT algorithm (which also applied soft thresholding in the denoising algorithm). The patch size was 3×3×1 voxels with a stride of 1 for all algorithms.

All denoised data and the original noisy data was fitted using an explicit three-peak “AMARES style” algorithm (equivalent to equation [9] with K=3; implementing bounds and manual starting values but no ‘prior-knowledge’ constraints) and optimized using the Scipy ‘curve_fit’ Trust Region Reflective algorithm (27).

For each noise level and algorithm, a spectral RMSE was calculated using the true noiseless data, and a fitted concentration RMSE was calculated from the input metabolite concentration maps (supporting information figure S1a).

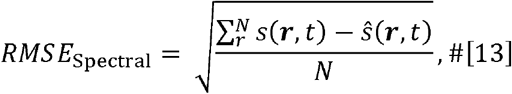

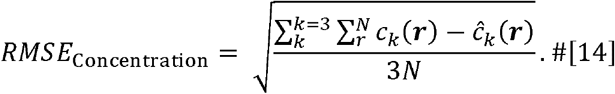

In addition, the voxel-wise concentration error was calculated, and the mean, standard deviation, and skewness of the resulting distribution calculated at each noise level. All metrics were calculated using all voxels and all MC repetitions (*N* = 8 × 8 × 50 × 3200 voxels).

## Results

### Denoising of uniform single-peak simulation

Monte Carlo measured frequency-domain standard deviation showed significant non-uniformity for all denoising methods, with the highest variance coinciding with regions of highest signal (Figure 1).

Qualitative evaluation of the proposed variance estimation method for the ST algorithm resulted in good agreement with the MC estimated variance (Figure 3) and the covariance (supporting figure S3).

**Figure 3.**
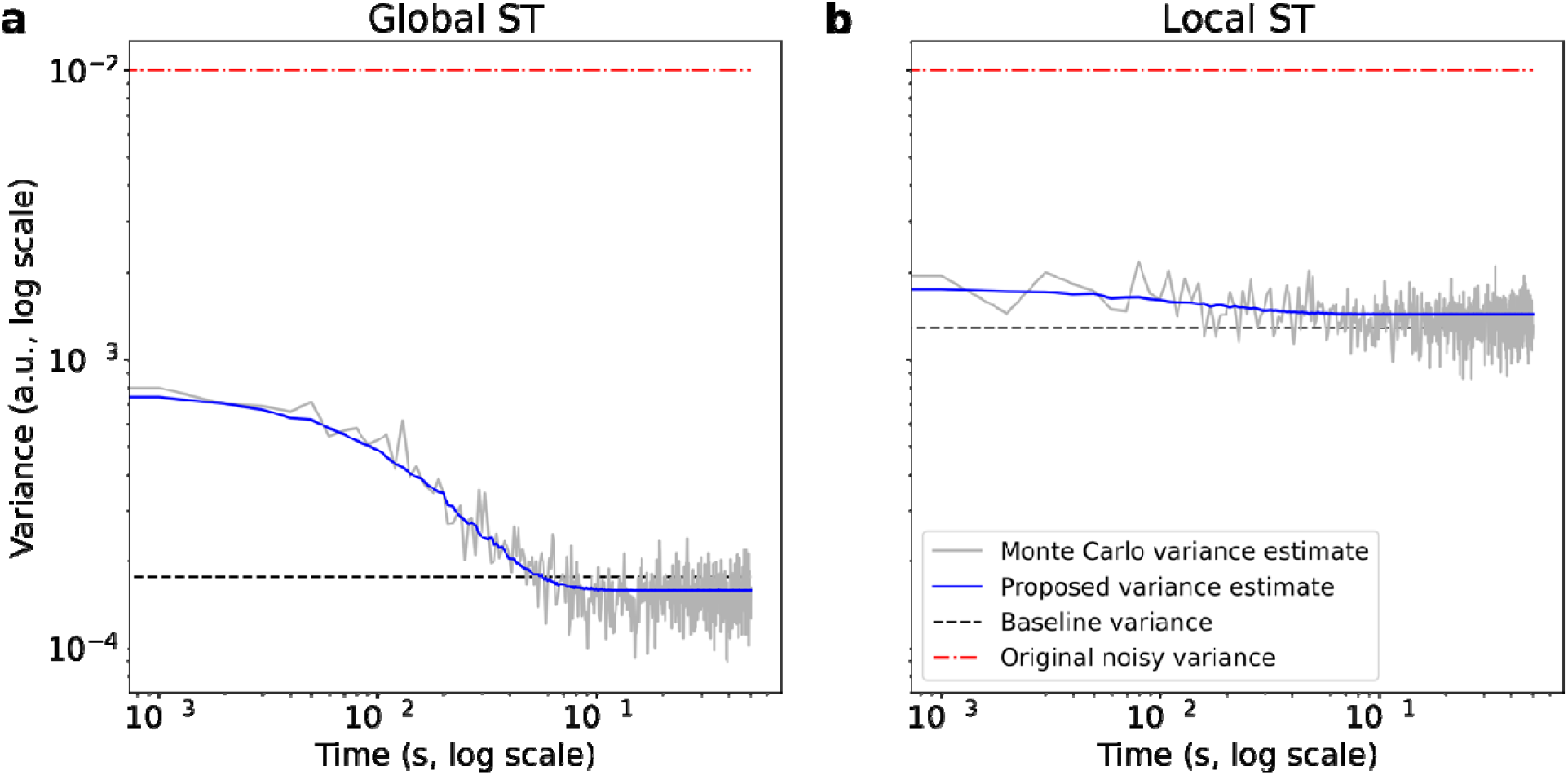
The variance estimated using the proposed method (blue) after global ST **(a)** and local ST **(b)** compared to the Monte Carlo variance estimate (grey). Also shown is the noise variance before denoising (red) and that which would be estimated from the denoised signal-free baseline (black).

For the single peak simulations, all algorithms except the LP algorithm decreased the uncertainty in fitted amplitude (Figure 4). The LP algorithm increased the uncertainty. Furthermore, the conventional uncertainty significantly under-estimated actual uncertainty for all algorithms (‘Fit’ vs ‘MC’ in Figure 4). The fitting algorithm estimated the conventional uncertainty for the Global ST method is equal to the uncertainty predicted for the spatial average of all voxels (vertical dashed line). The bootstrap method was able to accurately estimate the actual uncertainty (‘Bootstrap’ in Figure 4). These results were consistent across all noise levels examined (Supporting Figure S4).

**Figure 4.**
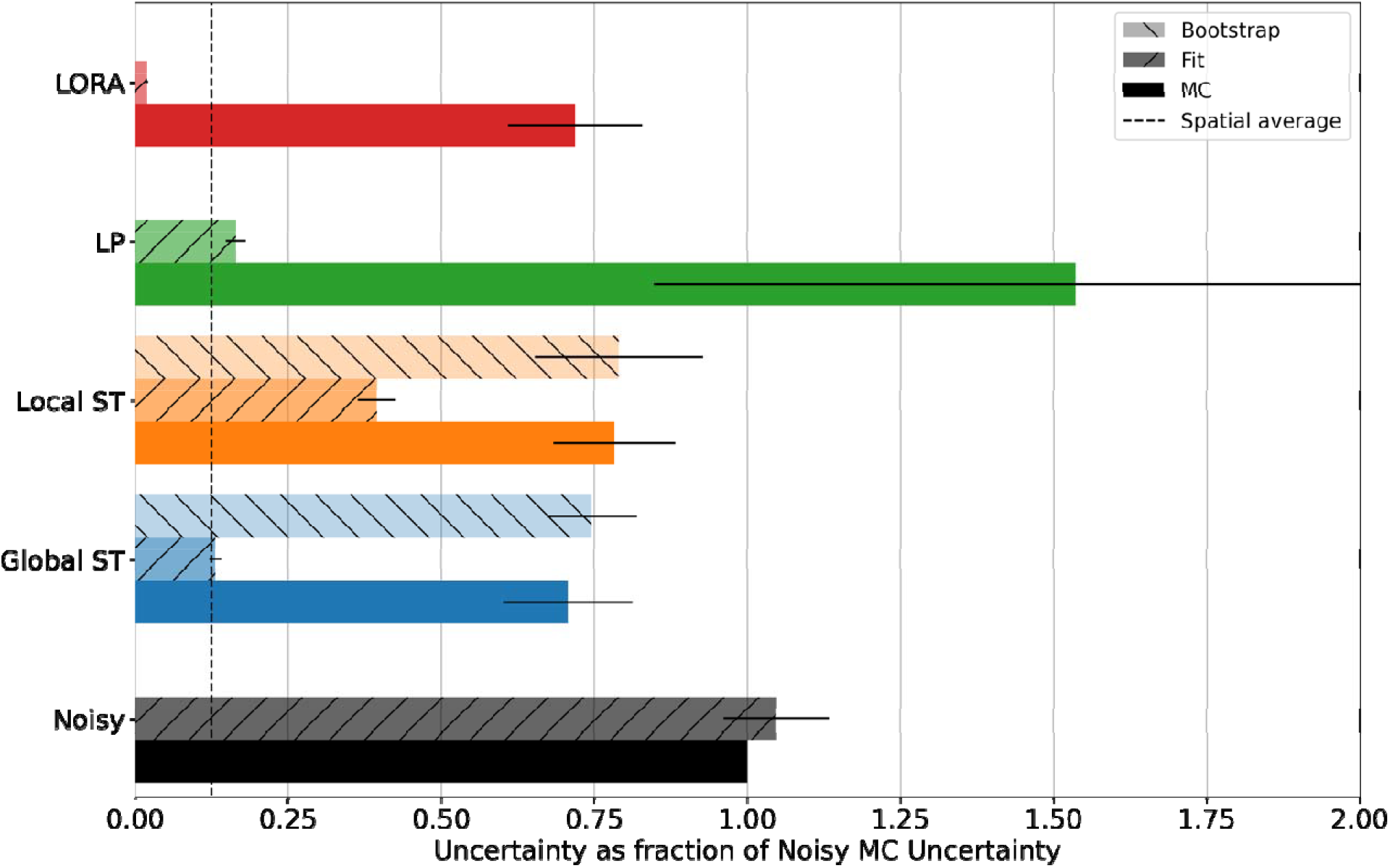
Single peak fitted amplitude uncertainty (standard deviation) relative to ‘Noisy’ for each denoising algorithm. Data shows the Monte Carlo estimated actual uncertainty (‘MC’), the conventional uncertainty (‘Fit’), and the Bootstrap estimated actual uncertainty (‘BS’), as mean and standard deviation across all simulated noise levels. The LP algorithm (green) increases uncertainty, all others decrease uncertainty. The conventionally estimated uncertainty is not accurate and for the Global ST case is equal to that which would be estimated for the spatial average of all voxels. The proposed bootstrapping method provides an accurate estimate of the uncertainty for both the local and global ST algorithms.

### Evaluation of denoising methods in simulated ^1^H-MRSI

For the 9-minute synthetic MRSI data all denoising algorithms, except LP, reduced the actual uncertainty (Figure 5a). Furthermore, in all cases the estimated conventional uncertainty was lower than the MC measured actual uncertainty, though that for the local ST method only underestimated by 17% (vs. 64% for global ST). Specifically, the mean (±SD) actual uncertainty ratio of the ‘high signal’ metabolites for the global ST, local ST, LP, global LORA, and local LORA methods were 0.67±0.20, 0.56±0.15, 1.07±0.20, 0.68±0.21, 0.57±0.15. The values were 0.57±0.42, 0.57±0.34, 1.22±2.40, 0.58±0.38, 0.65±2.55 for ‘all unique’ metabolites (Supporting Figure S5). The median normalized RMSE showed the same pattern (Figure 5b), with RMSE of 0.056, 0.050, 0.068, 0.057, 0.050 (noisy = 0.062).

**Figure 5.**
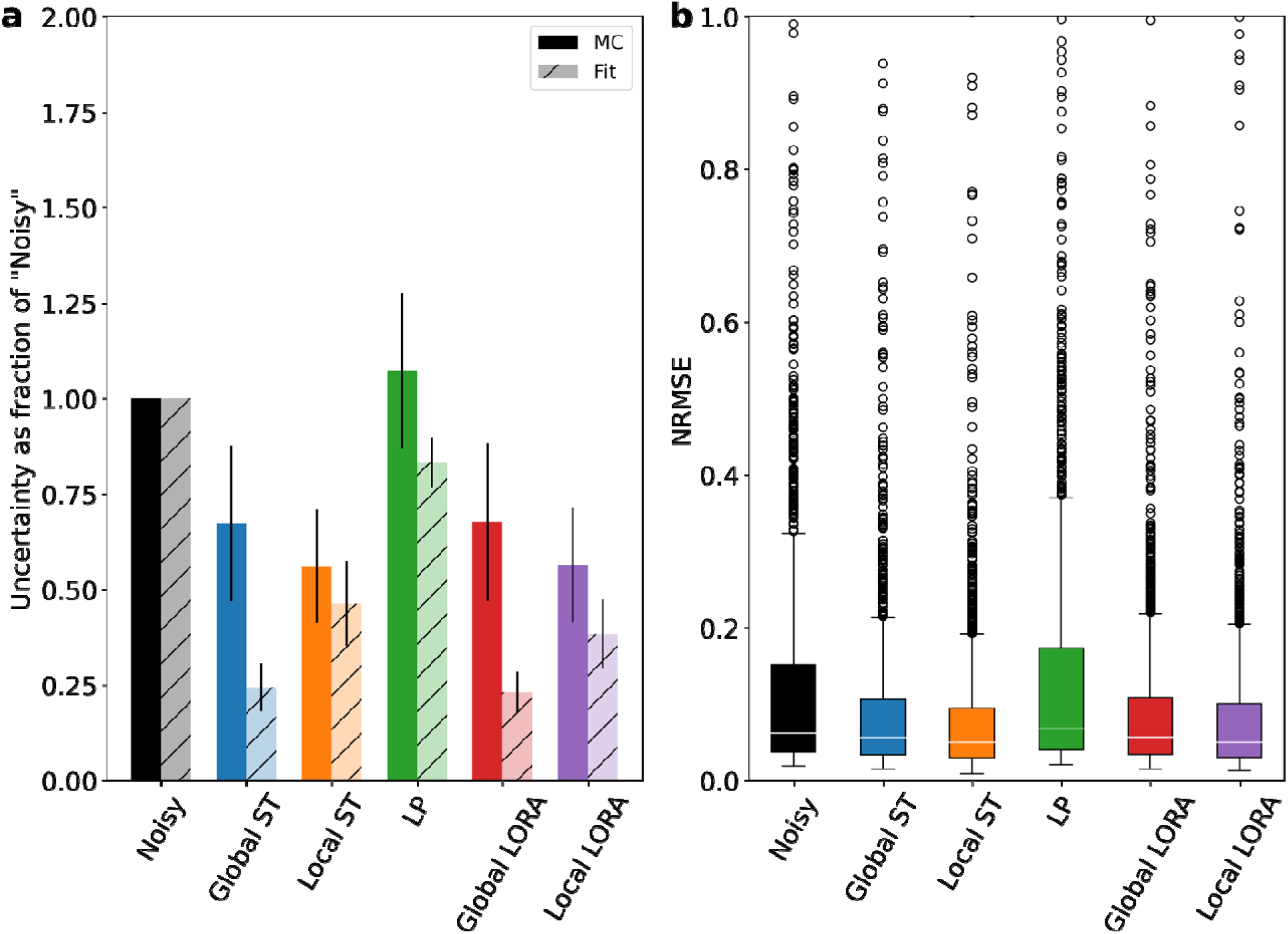
**a** Mean uncertainty of the fitted concentrations of the ‘high signal’ metabolites expressed as a ratio to the original noisy data. Actual uncertainty measured by MC simulation is compared with that conventionally estimated by the FSL-MRS fitting algorithm. **b** Normalized RMSE of the noisy and denoised data comparing the fitted concentrations of the ‘high signal’ metabolites to that of the noiseless synthetic data.

Qualitative evaluation of the variance estimated by the proposed method against that measured by MC simulation showed that the proposed method slightly underestimated the variance, but predicted its non-uniform time (or spectral)-dependence, as well as capturing features of the covariance structure (Supporting figures S6 & S7).

The local and global ST methods were assessed at all five noise levels. Both methods reduced the actual uncertainty in metabolite concentrations at all noise levels (Figure 6 and S8), with the local ST algorithm outperforming the global algorithm, and achieving a reduction of more than half at the highest noise level.

**Figure 6.**
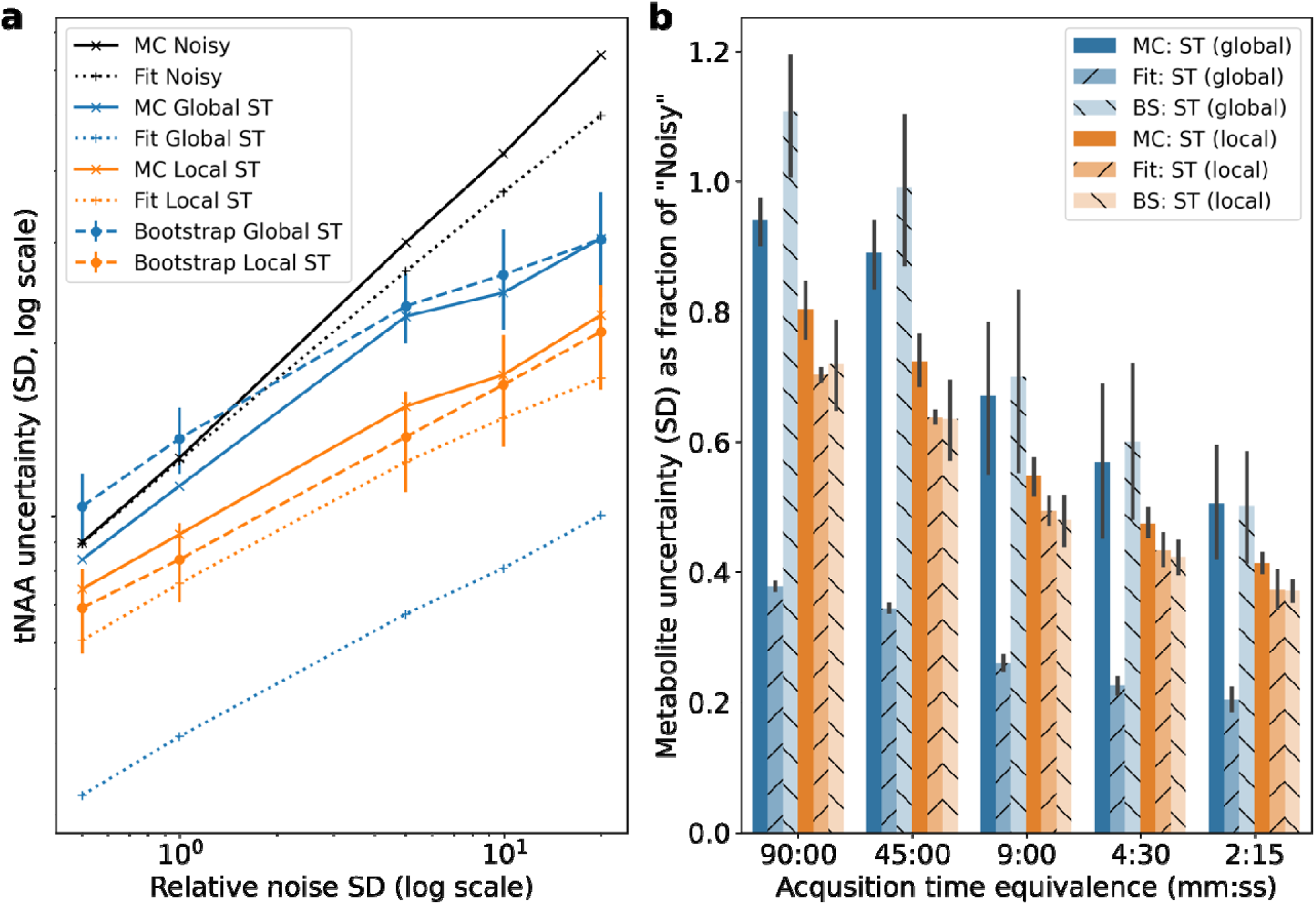
Estimated uncertainty at different noise levels by Monte Carlo simulation, FSL-MRS fitting, and bootstrap fitting for a single combined resonance **(a**; NAA+NAAG), and all ‘high signal’ metabolites **(b)**.

The bootstrapping uncertainty accurately predicted the actual uncertainty of the global ST method at all but the lowest noise levels, where it over estimated (bootstrapping = 107% of the actual uncertainty, conventionally estimated = 41%). For the local ST method, the bootstrapping method was as accurate as the conventional estimate: bootstrapping = 90% of the actual uncertainty, conventional = 90% of the actual uncertainty. Figure 6b illustrates this for each noise level for the ‘high signal’ metabolites. See supporting Figure S8 for the ‘all unique’ metabolites.

### Reproducibility of ^1^H-MRSI

In the in vivo data metabolite concertation RMSE was reduced by all ST and LORA methods, except for the highest signal metabolites: tNAA, tCr, tCho (Figure 7a). Although RMSE was lowered for these metabolites by the local patch-wise denoising implementations. The LP algorithm raised the RMSE in nearly all cases, and the ST algorithm outperformed the LORA algorithm.

**Figure 7.**
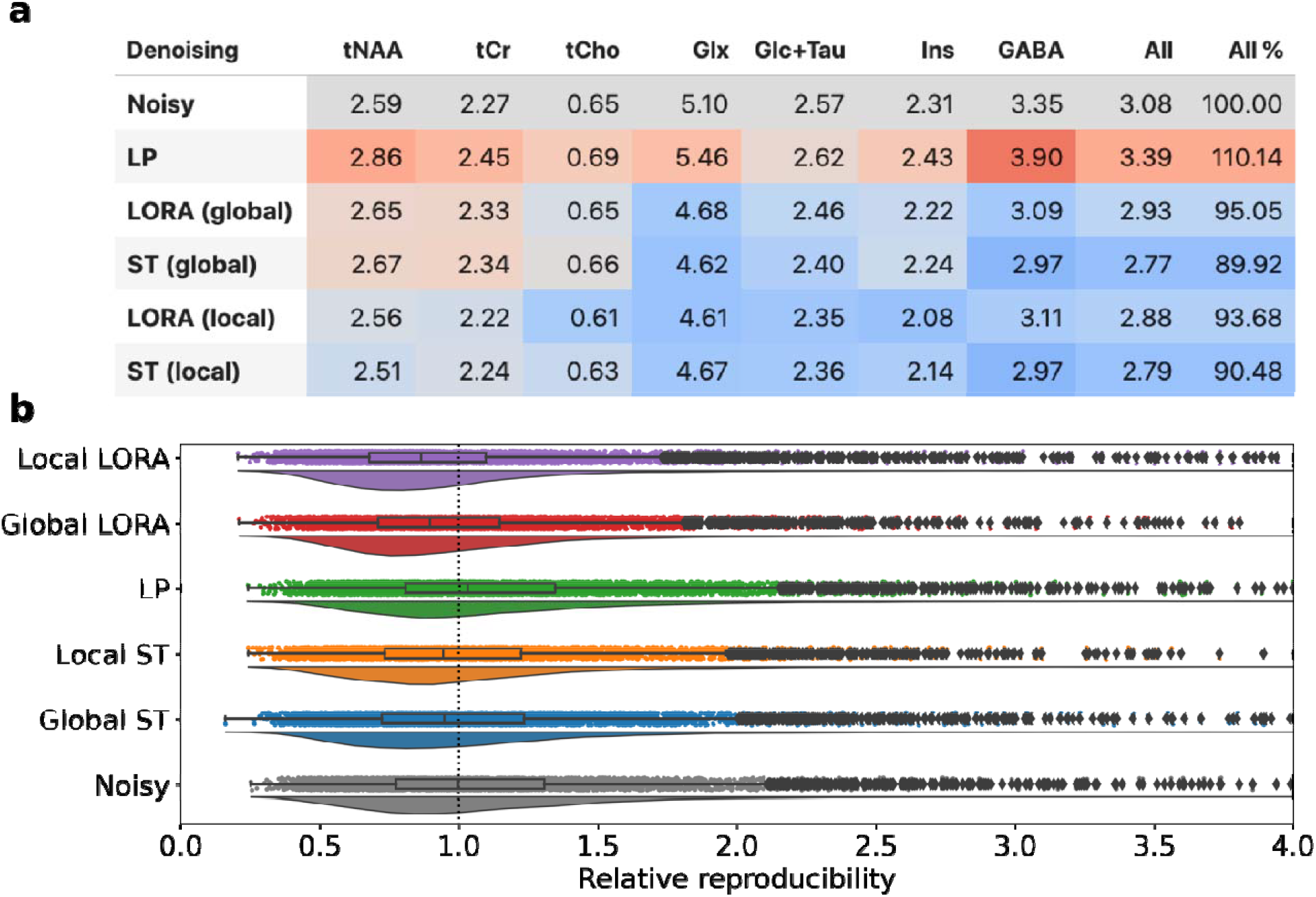
**a** RMSE for high signal metabolites in denoised 4.5-minute scans compared to the 45-minute equivalent average. Blue shading indicates values lower than the noisy baseline; red indicates values higher. **b** Rainfall plots of the relative reproducibility (standard deviation of the ten 4.5-minute scans) for the ‘high signal’ metabolites.

Relative reproducibility (Figure 7b) showed similar results with median (±SD) ratios of 0.95±0.53, 0.94±0.53, 1.03±0.59, 0.90±0.48, 0.86±0.52 for global ST, local ST, LP, global LORA, and local LORA respectively.

### Bias and rank selection in simulated data

Figure 8a & 8b shows the spectral RMSE and concentration RMSE respectively. SURE-optimized SVHT produced the lowest Spectral RMSE, whilst MP produced a consistently low Concentration RMSE. At high noise levels (indicated by vertical dotted line in Figure 8b) SURE SVT and the fixed R = 2 algorithm achieved lower concentration RMSE than MP.

**Figure 8.**
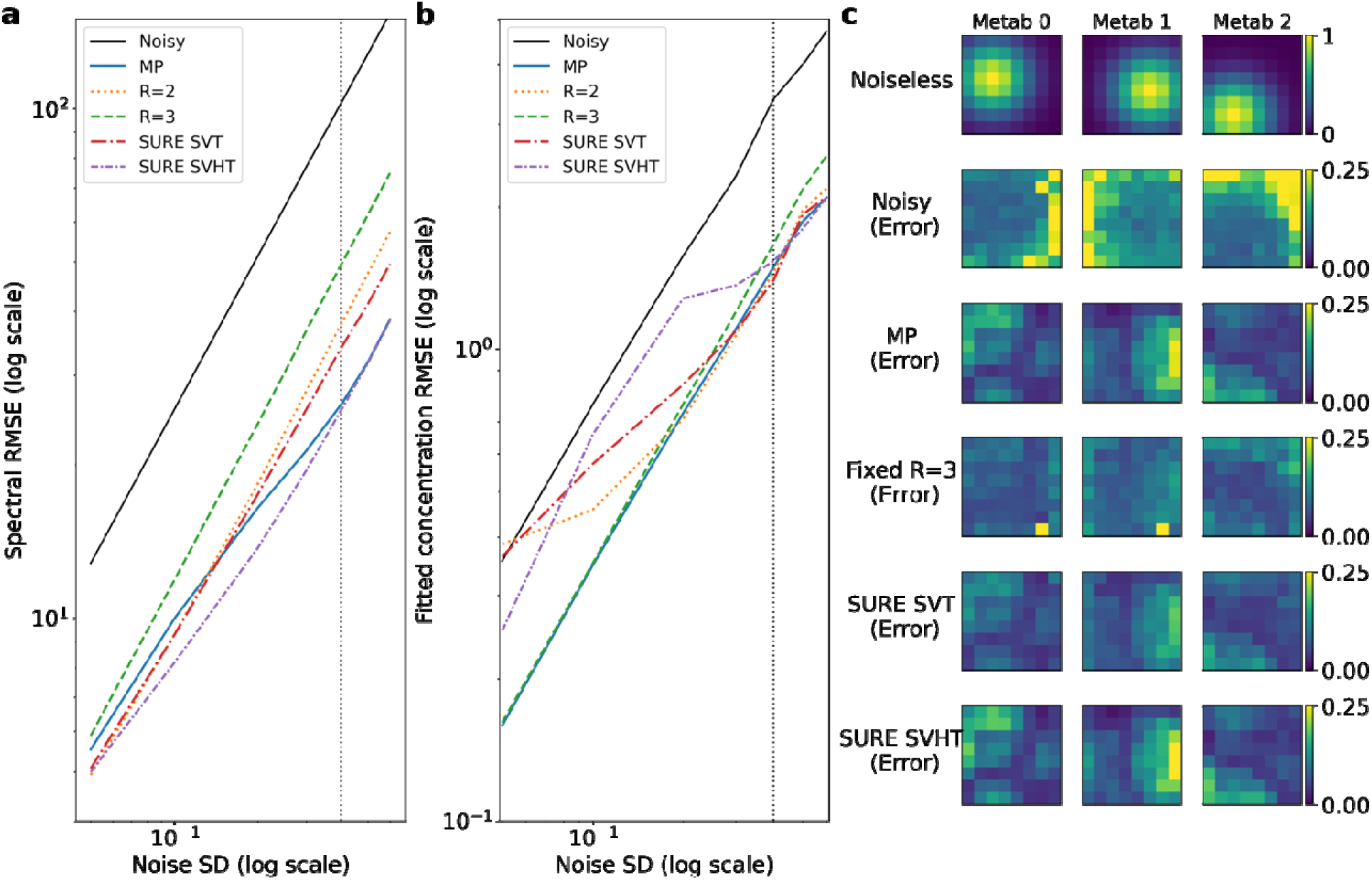
Spectral RMSE **(a)** and concentration RMSE **(b)** of the five different thresholding approaches of the simulated rank = 3 data, compared to the noisy data. Panel **c** shows the noiseless concentration maps for each of the three ‘metabolites’ and the absolute error for each method at a single noise level (dotted vertical line in **a** & **b**). High bias is observed for SURE SVHT and MP, with less seen in SURE SVT and R=3. However, the fixed R=3 shows higher average error.

Voxel-wise error also showed that SURE SVT had lower mean error, skew, and equivalent standard deviation to MP at high noise levels, despite relatively poor performance at low noise levels (Figure 9d-f). MP thresholding has high bias and skew at the highest noise levels but outperforms SURE SVHT at most points as measured by concentration RMSE.

**Figure 9.**
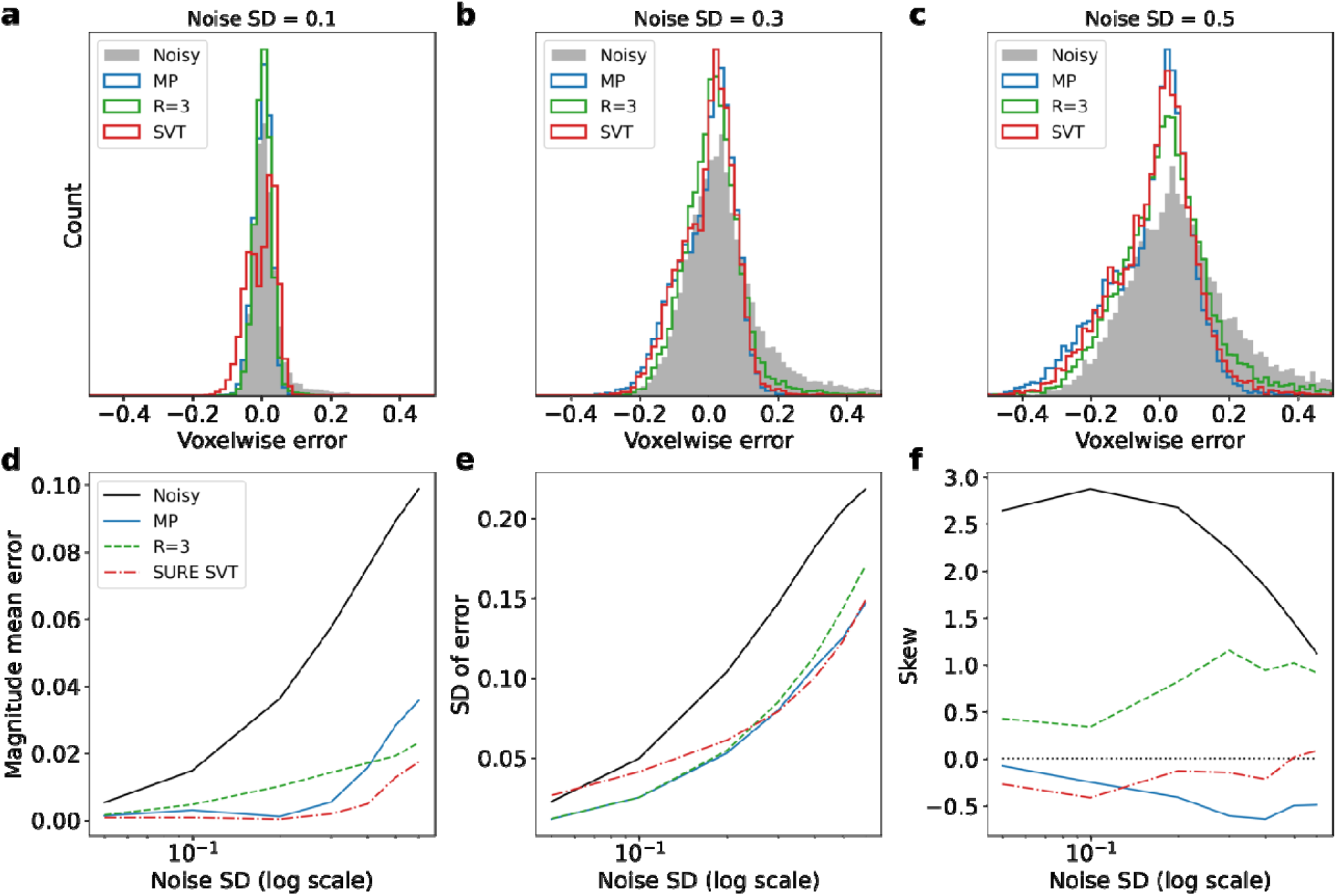
**a-c** Voxel-wise error distributions for three methods at three different noise levels. **d-f** Magnitude mean error, standard deviation, and skew of the distributions formed by the voxel-wise error.

## Discussion and Conclusion

This work has demonstrated that low-rank denoising based on spatio-temporal separability can reduce uncertainty of estimated metabolite concentrations in both synthetic and in vivo ^1^H-MRSI data. However, the reduction in uncertainty is typically not as great as is apparent, because the non-uniform residual variance is under-estimated using conventional baseline noise inspection methods. Furthermore, denoising based on spectral linear predictability confers no benefit, as seen with increased uncertainty after LP denoising in all tests, and the similar performance of LORA when compared to ST denoising alone. In simulated ^1^H-MRSI data it is also apparent that patch based (local) methods perform better than global methods. Local methods provided no improvement in the single-peak case, or for the in vivo data, though the latter may be masked by inherent inter-scan variability which cannot be removed by denoising, and optimal patch sizes are likely data-dependent. It should be noted that LORA might still be advantageous for situations where higher apparent denoising is desirable, for example generating basis spectra.

The proposed uncertainty estimation method (via bootstrapping) was found to be highly accurate in the case of a single peak. Though not shown in this work, the same was found to be the case when extended to simple multi-peak simulation. In simulated 1H-MRSI data denoised by global ST the bootstrap uncertainty was much closer to the MC-measured actual uncertainty than the conventional uncertainty estimate (107% vs. 41%). For the local ST method, both the uncertainty estimated conventionally and the proposed method were relatively accurate (both 90% of actual). This may be because the repeated averaging of patches removes the non-uniform variance and covariance that is apparent in the global case. This work has not assessed how many overlapping patches are needed to achieve this, nor its dependence on patch stride or size. This may become important for data with fewer dimensions, e.g. one dimensional dynamic (functional or diffusion weighted) MRS.

If more accurate covariance estimation was available, the bootstrapping uncertainty estimation could be improved. In simulated ^1^H-MRSI data the variance and covariance estimation method underestimated the denoised data variance and covariance (supporting Figures S6 & S7). Our estimate of the off-diagonal covariance elements accurately captures the structure, but not the magnitude in a range of data (Supporting Figure S7 and supporting text “Covariance approximation”). Chen et al note that the variance estimation (Equation 6) deteriorates at low SNR and has a dependence on condition number. Bootstrap fitting of MRSI data is slow and only 40 repetitions were used here in the simulated H-MRSI data, more might also lead to higher accuracy.

In this work we introduce the use of SURE and soft thresholding to the denoising of MRSI data. Whilst SURE selection of a hard threshold minimizes spectral RMSE, it does not also minimize the RMSE of the fitted metabolite concentrations. Both SURE and MP automated threshold selection lead to biased data in low SNR cases (as the rank threshold tends to 1 and signal variance is lost). The lower bias and skewness of the SURE SVT (soft thresholding) at high noise levels indicates that it or other thresholding functions may be optimal for low SNR regimes (or where trying to detect small signal fluctuations), such as phosphorus-31 MRS or diffusion weighted MRS (9,28). In fact, recent work investigating optimal matrix denoising (29) highlights the increased performance of soft-thresholding in lower SNR regimes, and also proposes optimized singular value shrinkage functions that out-perform hard or soft-thresholding in all SNR regimes. Evaluation of this parameter-free optimized singular value shrinkage method would certainly be an interesting direction for future work.

In this work we have not assessed the impact of algorithm parameters: undoubtably patch size and stride for the local methods are important. Currently there is no automated way to select these. In addition, it is difficult to assess the true uncertainty reduction in in vivo data. Here we have demonstrated modest (10%) decreases in inter-scan variability, but the true amount might be masked by inter-scan variability not caused by thermal noise e.g., physiological noise, or scanner drift.

The denoising tools used in this work have been made available for use as the Python package ‘mrs_denoising_tools’ and operate on four dimensional MRSI data stored in the NIfTI format.

In conclusion, low-rank spectroscopic denoising methods based on spatio-temporal (or dynamic-temporal) separability do reduce uncertainty in MRS(I) data. However, thorough assessment of the method and use case should be made. It is important to select the right thresholding method in low SNR cases, and assessment simply by SNR measured from residual baseline noise is insufficient given the possibility of non-uniform variance.

## Data availability statement

The source code and data used to generate the results presented in this manuscript can be found at https://git.fmrib.ox.ac.uk/wclarke/low-rank-denoising-uncertainty (#8612a5ce52b254f987aae8291e74af7c19d9a685).

The specific MRSI denoising tools have been released as a Python package “mrs-denoising-tools” available from the Pypi and Conda package managers. The source code is available from https://git.fmrib.ox.ac.uk/wclarke/low-rank-denoising-tools. This work was created using version 0.0.2 of mrs-denoising-tools.

Fitting was performed using version 1.1.2 of FSL-MRS, which is also available from the Conda package manager and https://git.fmrib.ox.ac.uk/fsl/fsl_mrs.

## Acknowledgements

This research was funded by the EPSRC (EP/T013133/1), the Royal Academy of Engineering (RF201617\16\23), and the Wellcome Trust and the Royal Society (102584/Z/13/Z).

The Wellcome Centre for Integrative Neuroimaging is supported by core funding from the Wellcome Trust (203139/Z/16/Z).

We thank Adam Steel and Charlotte Stagg for providing the in vivo data.

