## Supporting Information for "Uncertainty in denoising of MRSI using low-rank methods"

Supporting information: Uncertainty in denoising of MRSI using low rank methods

### Covariance approximation

In this work Equation 7 is used to approximate the covariance of the time domain data after ST denoising has been applied. This section contains an assessment of the accuracy of this approximation. The assessment is carried out with five different test cases, at two noise levels. For each case, the Equation 7-estimated covariance is compared with the Monte Carlo measured covariance of a single voxel. The test cases shown here are:

1. A randomly generated low-rank matrix.
2. 128 voxels of a single-on resonance peak as generated for the section “Denoising of uniform single-peak simulation”
3. 128 voxels of a single-off resonance peak as generated #2 with a 100 Hz shift.
4. 64 voxels of explicitly rank=3 data as generated for the “Bias and rank selection in simulated data”. Noiseless data is shown in Supporting figure S1.
5. 64 voxels with 20 different peaks, resulting in a matrix of approximately rank=20.

Error was quantified as the element-wise difference normalised by the maximum of the MC estimated covariance.

$$Error= 100*\frac{\left| cov_{\mathrm{est}}-cov_{\mathrm{MC}} \right|}{max\left| cov_{\mathrm{MC}} \right|}$$

Across all test cases, in the high noise case the highest mean error was 3% and the maximum error was 18%. In the low noise case the maximum mean error was 3% and the maximum error was 17%. In all cases the approximation captured the broad structure of the covariance accurately. Qualitative comparison of the accuracy of this method is also shown for data used in the main analysis in Figures S3 (single peak) and S7 (simulated 1H-MRSI).

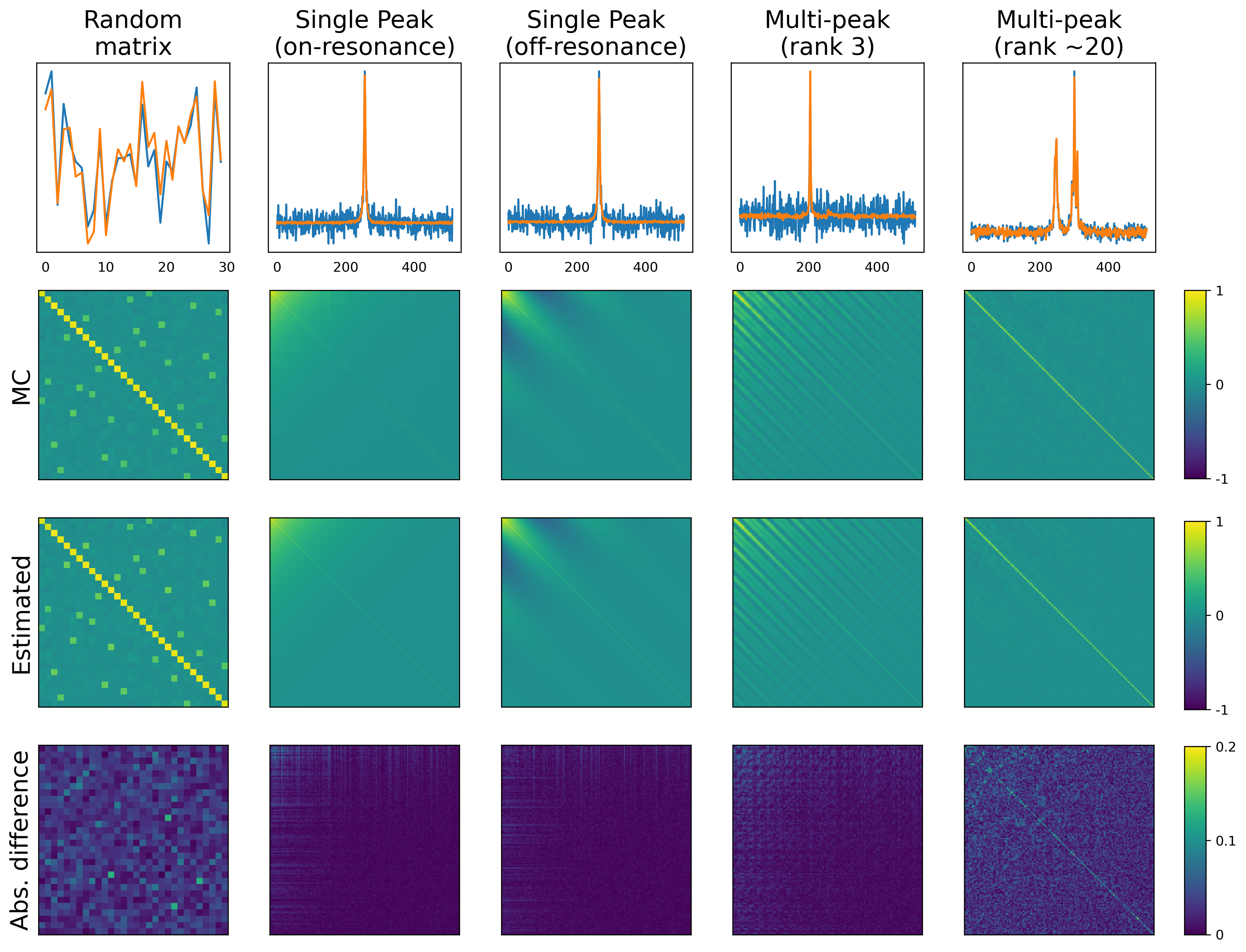

*Comparison of the Monte Carlo calculated time-domain covariance with that estimated using Equation 7 for the five test cases. The top row shows the noisy and denoised data from a single voxel. The bottom row shows the absolute difference between the MC and Estimated covariances (scale 0-20%).*

### Stein's Unbiased Risk Estimate (SURE)

The SURE equations for SVT, from Candès et al (14), and SVHT, adapted from Ulfarsson and Solo (15), are reproduced in full here.

$$SURE\left( SVT_{\lambda} \right)= -2mn\tau^{2}+\sum_{i=1}^{\min\left( m,n \right)} \min\left( \lambda^{2}, \sigma_{i}^{2} \right)+2\tau^{2}\mathrm{div}\left( SVT_{\lambda}\left( \boldsymbol{M} \right) \right),$$

$$\mathrm{div}\left( SVT_{\lambda}\left( \boldsymbol{M} \right) \right)=\left( 2\left| m-n \right|+1 \right)\sum_{i=1}^{\min\left( m,n \right)} \left( 1-\frac{\lambda}{\sigma_{i}} \right)_{+}$$

$$+\sum_{i=1}^{\min\left( m,n \right)} I(\sigma_{i}>\lambda)+4\sum_{i\neq j,i,j=1}^{\min\left( m,n \right)} \frac{\sigma_{i}\left( \sigma_{i}-\lambda\right)_{+}}{\sigma_{i}^{2}-\sigma_{j}^{2}},$$

where *m* and *n* are the dimensions of matrix ***M***, $\lambda$ is the threshold, $\sigma_{i}$ is the i^th^ singular value of ***M***, $\tau$ is the standard deviation of the i.i.d. noise, and $x_{+}=max(x,0)$.

$$SURE\left( SV{HT}_{R} \right)= -2mn\tau^{2}+\sum_{i=1}^{R} \sigma_{i}^{2}+4\tau^{2}\mathrm{div}\left( SVHT_{R}\left( \boldsymbol{M} \right) \right),$$

$$\mathrm{div}\left( SV{HT}_{R} \right)=R\left( m+n \right)-R^{2}+\sum_{i=1}^{R} \sum_{j=R}^{\min\left( m,n \right)} \frac{2\sigma_{j}^{2}}{\sigma_{i}^{2}-\sigma_{j}^{2}},$$

where $SV{HT}_{R}$ is implemented with a threshold of rank *R*.

SURE accurately predicts the MSE of the denoised spectral data from the singular values, matrix dimensions and i.i.d. noise variance of the input noisy data. An example of the SURE-predicted MSE in data identical to that shown in Supporting Figure S1 is shown below.

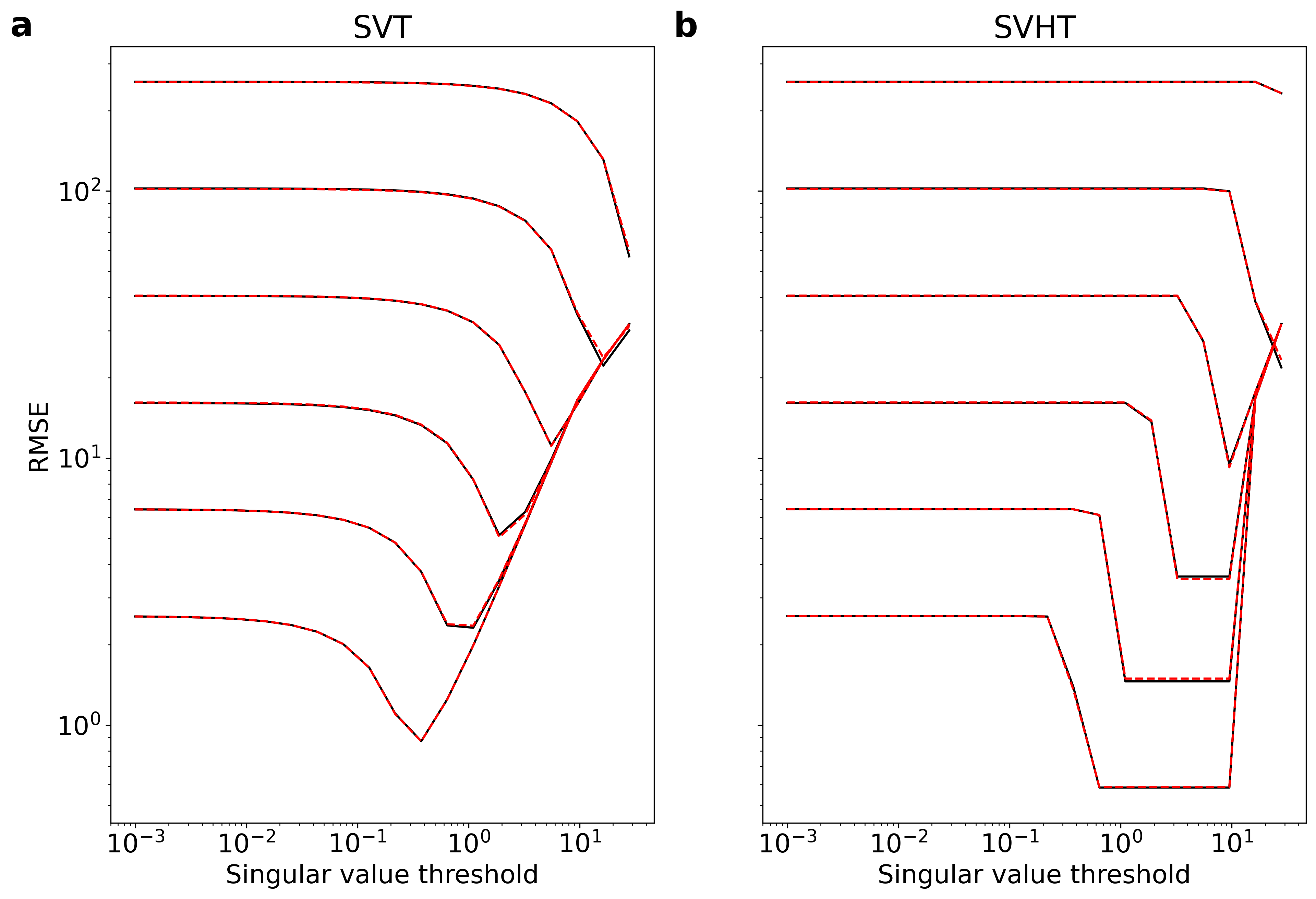

*SURE predicted RMSE (red dashed line) compared to numerical RMSE (black solid line) at six noise levels for:* ***a*** *soft thresholding,* ***b*** *hard thresholding. Noiseless input data is identical to that shown in Supporting Figure S1.*

### Supporting Figures

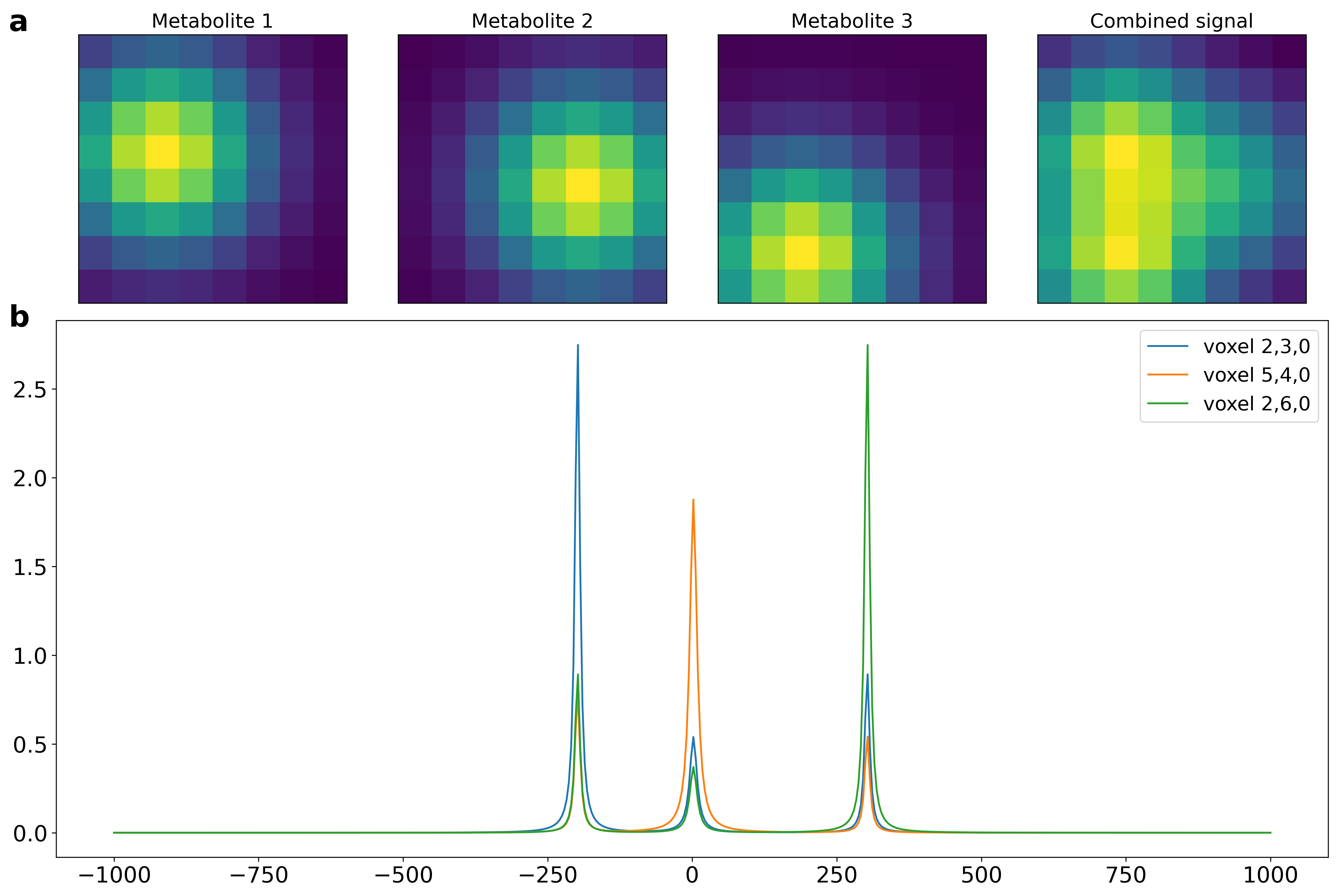

Supporting Figure S1: **a** ‘Metabolite’ maps for the three peak explicit rank-3 simulation data. **b** Spectra from three voxels of the same simulation data showing different relative ‘metabolite’ concentrations.

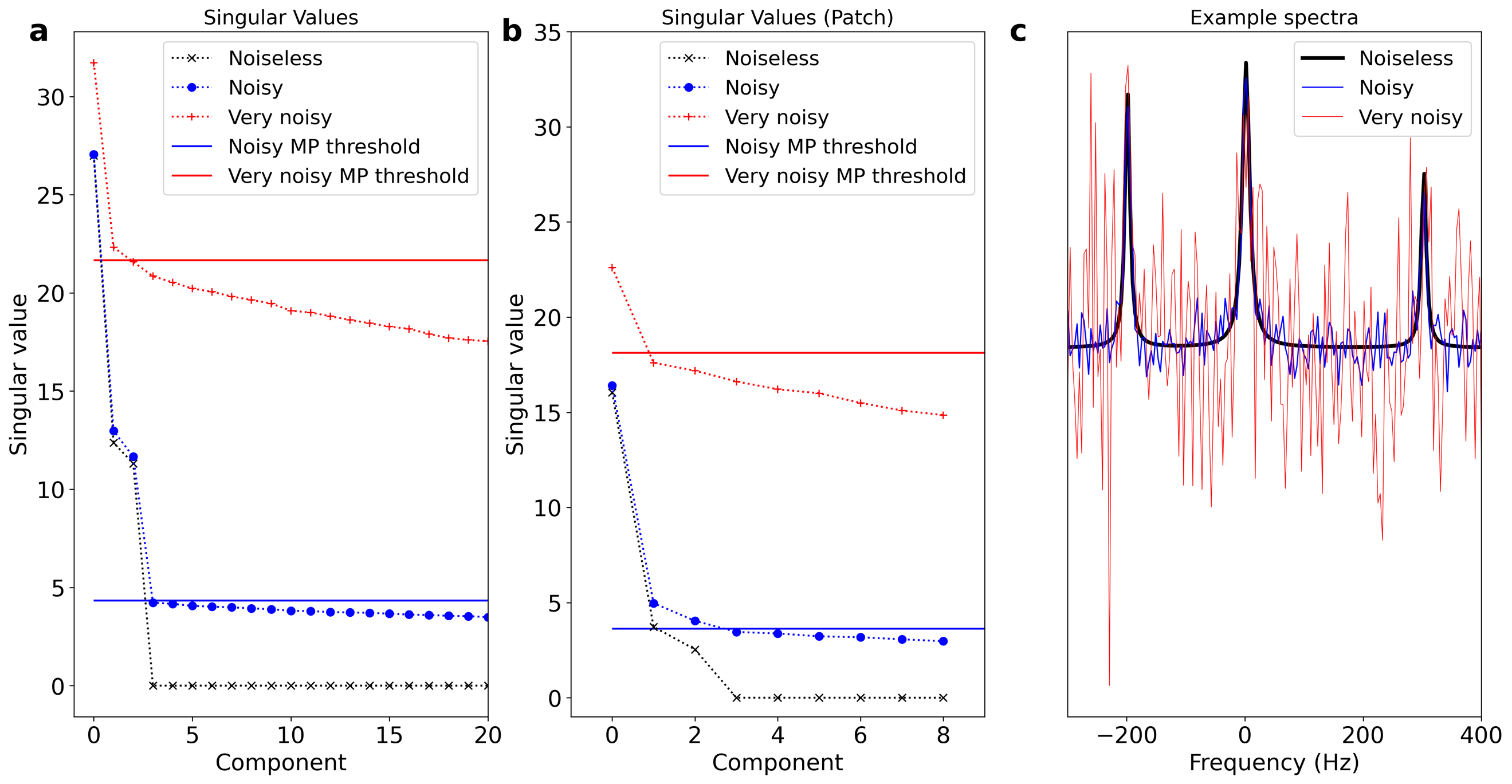

Supporting Figure S2: Underestimation of the rank of the data by the MP algorithm in noisy data. The solid horizontal lines show the estimated MP threshold in the noisy (blue) and very noisy (red) case. In the very noisy case, the rank is underestimated as 2 in the global case (**a**) and as 1 in the local case (**b**). Panel **c** shows example spectra and the relative noise levels.

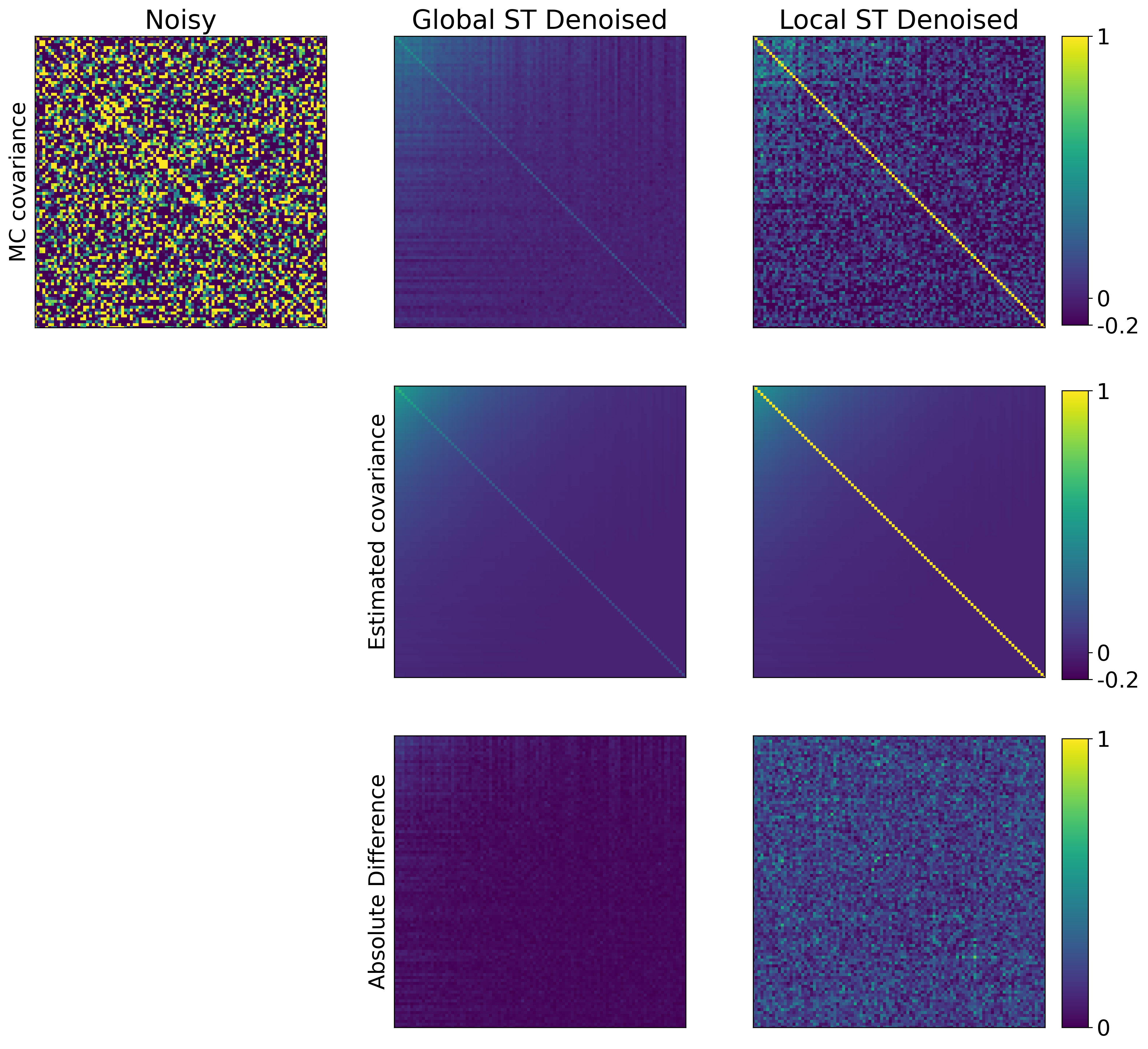

Supporting figure S3: Monte Carlo covariance and estimated covariance for the global and local ST denoised single peak data. There is close agreement between them. All plots are shown with the same arbitrary colour scale.

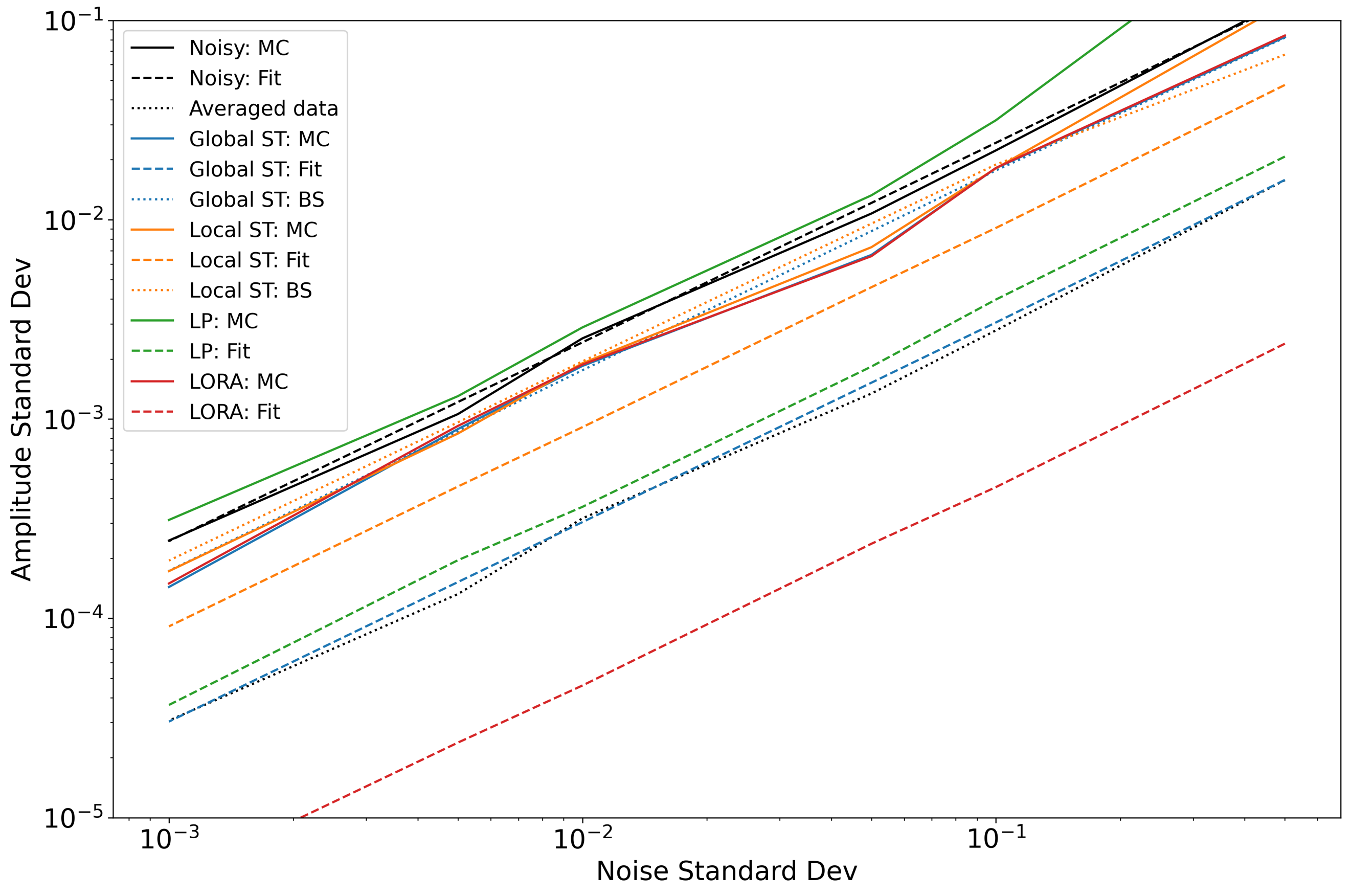

Supporting Figure S4: Uncertainty (standard deviation) of the single peak amplitude as a function of noise standard deviation. This plot shows the same data as Figure 4 but resolved across all input noise levels. Solid lines show the uncertainty estimated by Monte Carlo simulation, dashed lines by the fitting algorithm, and dotted lines by the proposed bootstrap process. The line ‘averaged data’ shows the uncertainty that would be achieved if all untreated (noisy) voxels were averaged before fitting.

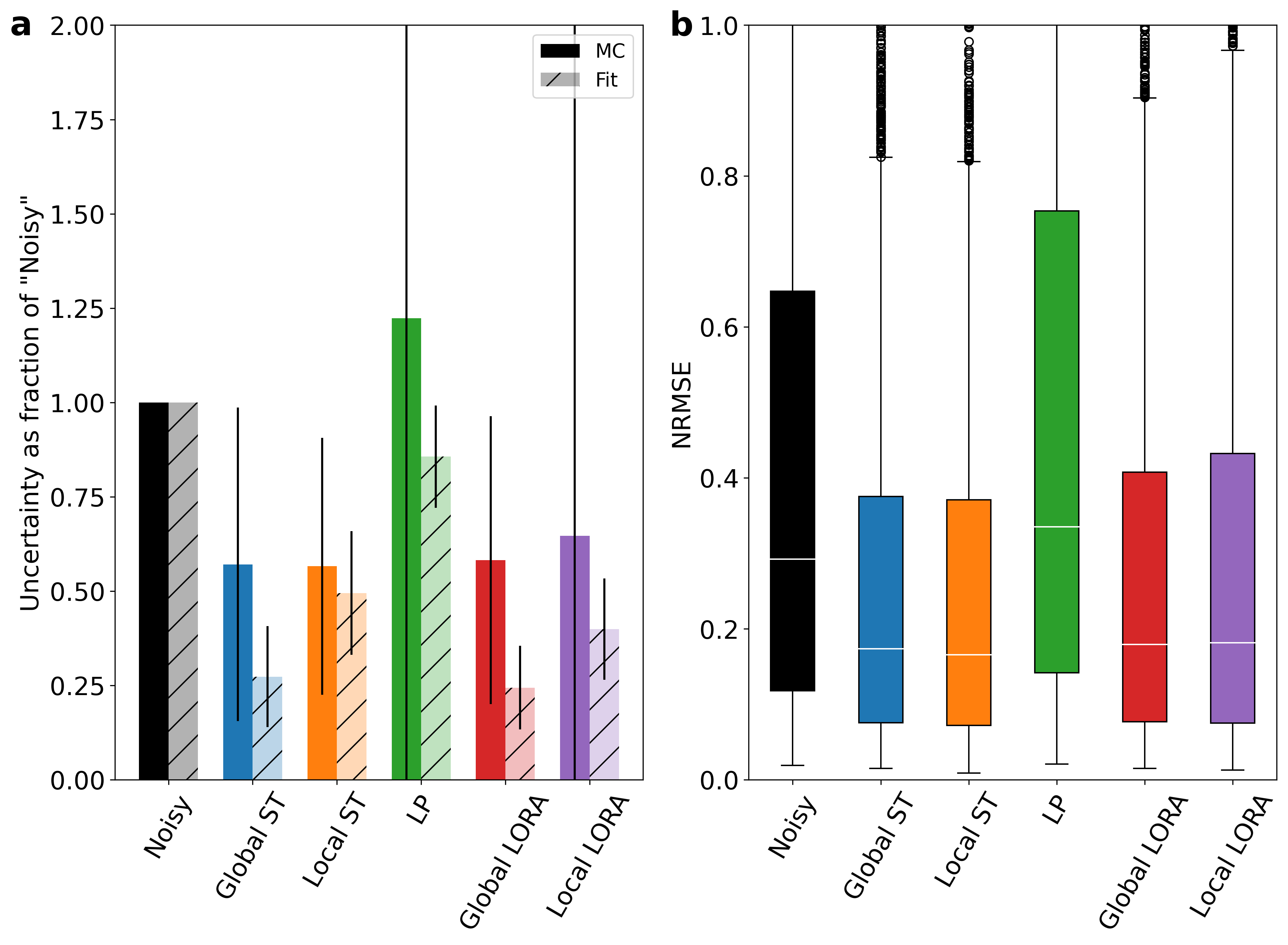

Supporting Figure S5: Equivalent to Figure 5 for the ‘all unique’ group of metabolites. **a** Mean uncertainty of the fitted concentrations of the ‘all unique’ metabolites expressed as a ratio to the original noisy data. Uncertainty measured by MC simulation is compared with that estimated by the FSL-MRS fitting algorithm. **b** Normalized RMSE of the noisy and denoised data comparing the fitted concentrations of the ‘high signal’ metabolites to that of the noiseless synthetic data.

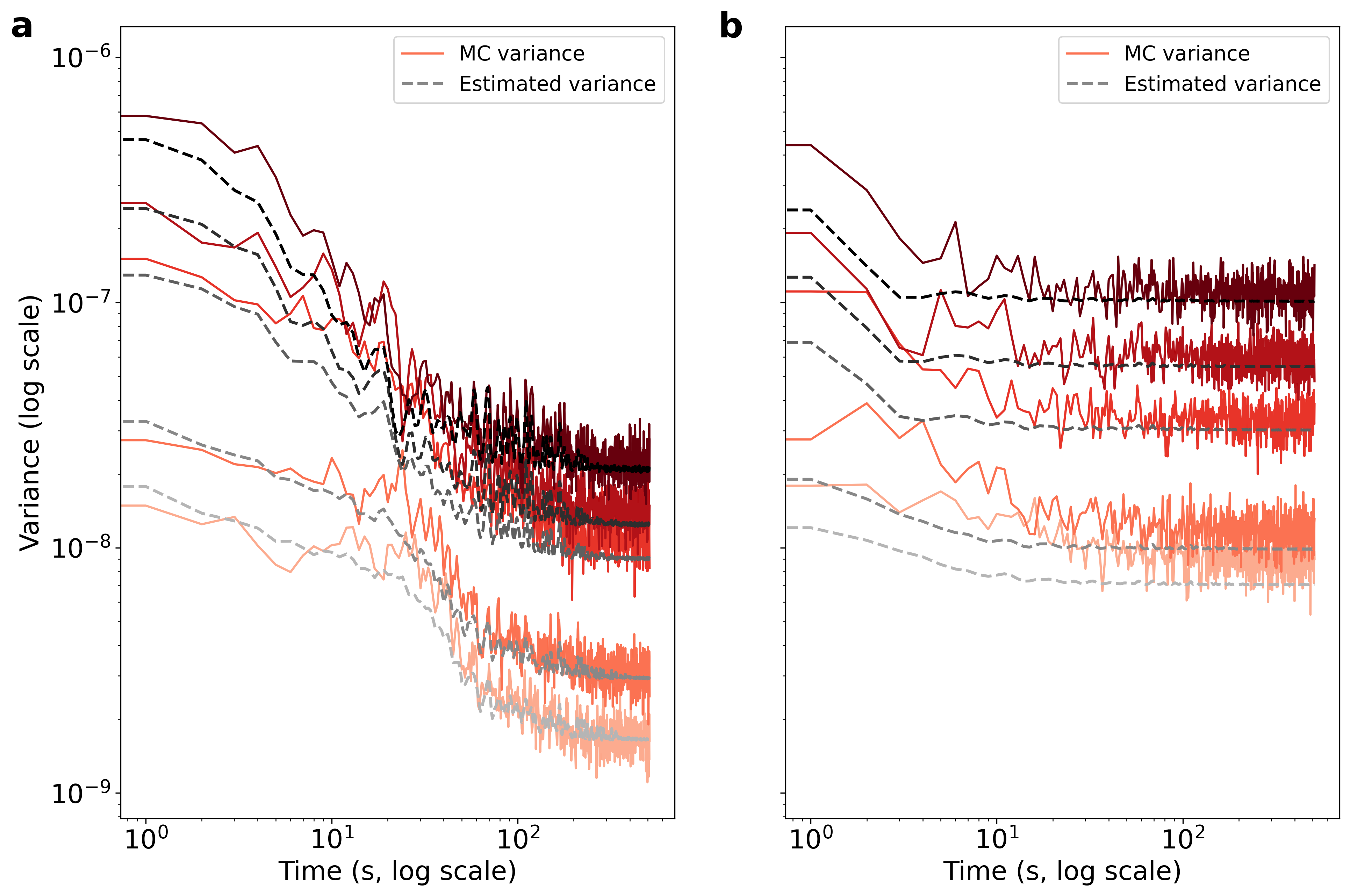

Supporting Figure S6: Estimated and MC measured variance for a single voxel of the simulated 1H-MRSI data at each of the five different noise levels. Panel **a** shows the variance for the global ST method and **b** shows that for the local ST method. The estimated variance captures the structure of the true (MC) variance, but globally underestimates the magnitude.

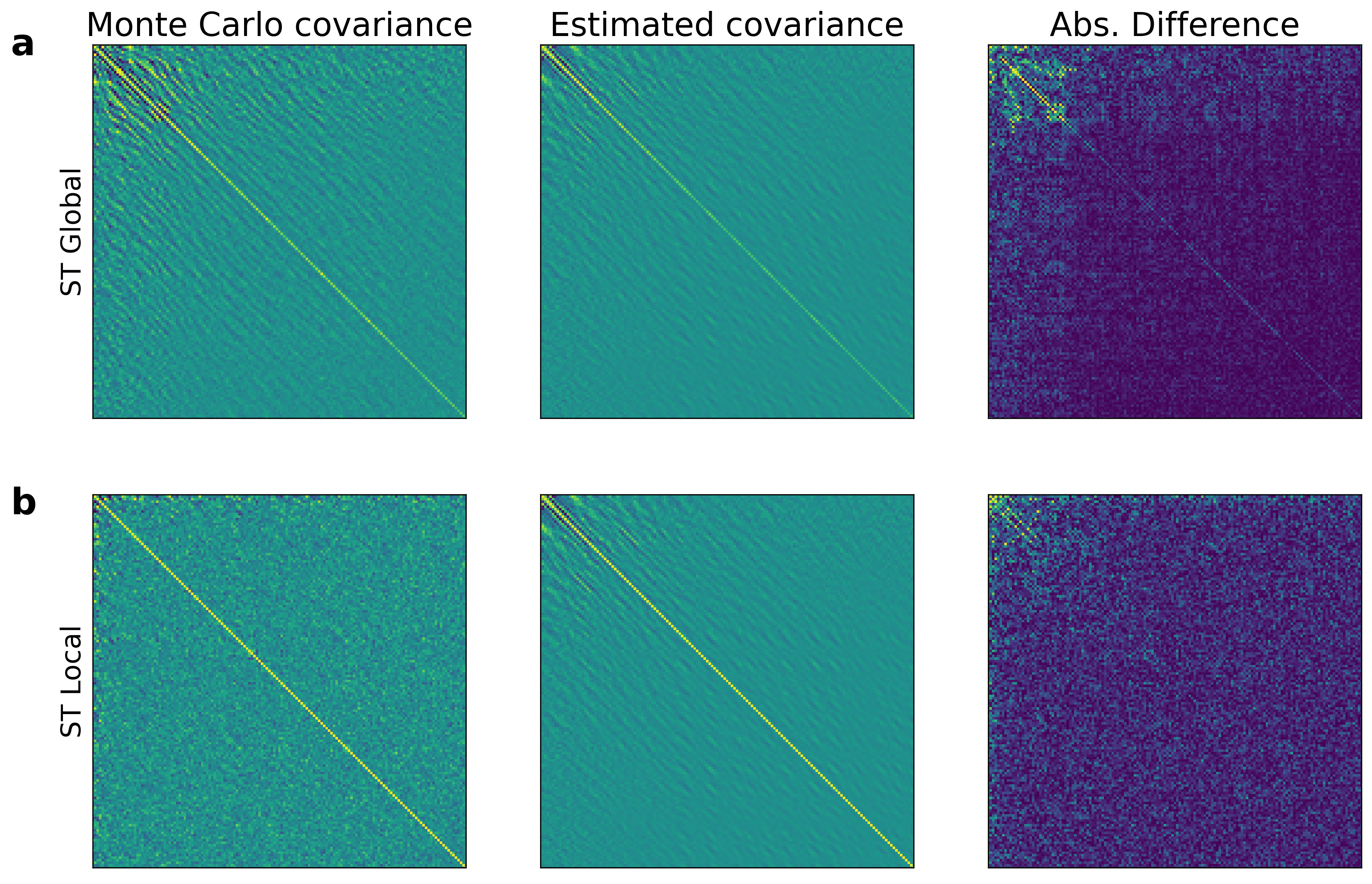

Supporting Figure S7: Covariance estimated using the proposed method and MC measured covariance for a single voxel of the simulated 1H-MRSI data at a single noise level (9-minute equivalent). Panel **a** shows the covariance for the global ST method and **b** shows that for the local ST method. The main diagonal slightly underestimates the variance (as in Figure S6), nor does the estimated covariance fully capture the complex covariance structure of the actual (MC) covariance in either case.

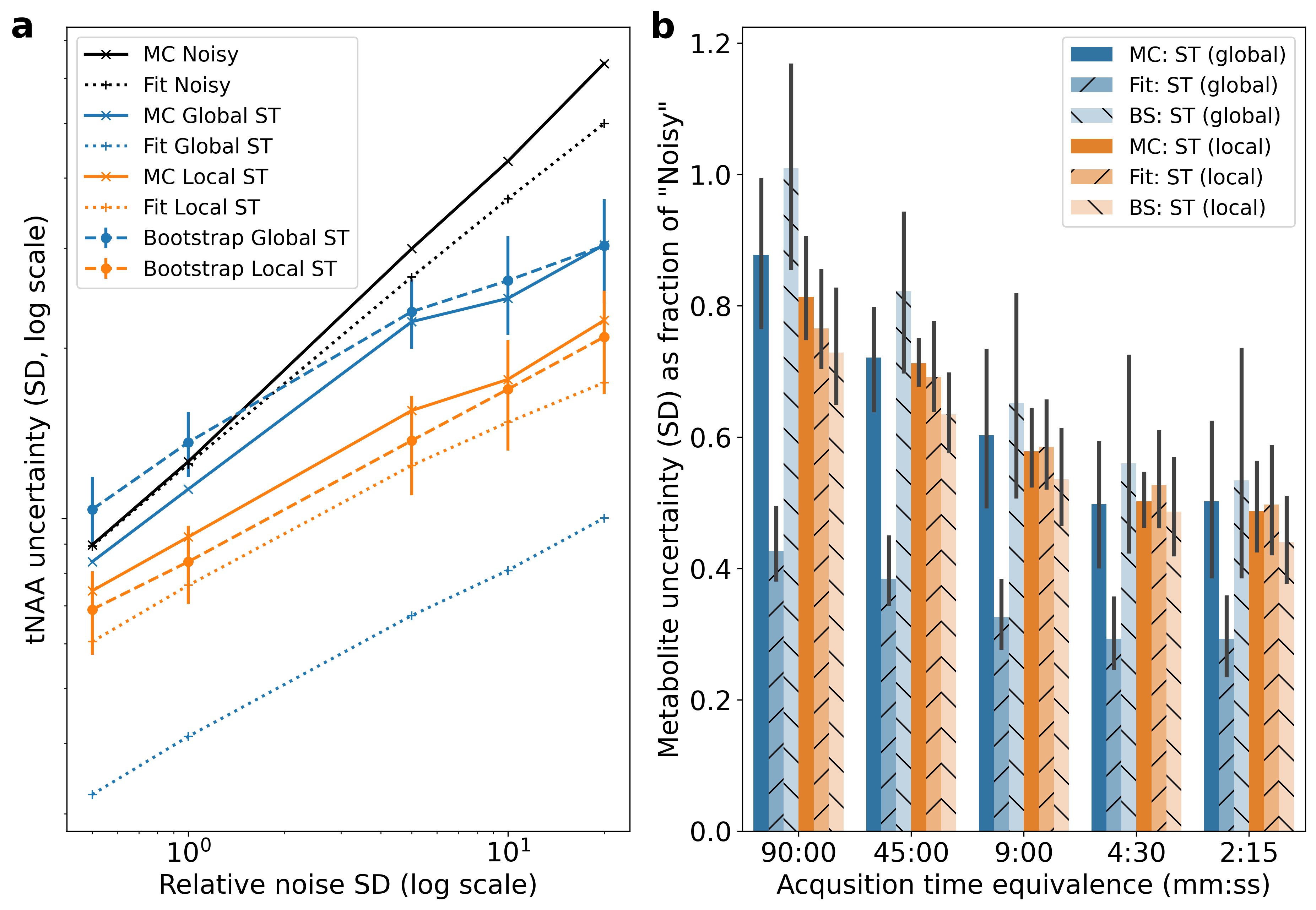

Supporting Figure S8: Estimated variance at different noise levels by Monte Carlo simulation, FSL-MRS fitting and bootstrap fitting for a single combined resonance (**a** NAA+NAAG) and ‘all unique’ metabolites (**b**). This plot is equivalent to figure 6 but for the different grouping of metabolites.
